# MaaD-TAD: an all-atom model of the duplicated PRAMEF/HNRNPCL chromatin domain built from sequence homology

**DOI:** 10.64898/2026.10.29.762793

**Authors:** Joy Ramielle L. Santos, Maya MacIsaac, Eun Bin Moon, A. Dean Befus, Marcelo Marcet-Palacios

## Abstract

In the nucleus, the placement of genes in the 3D fold of DNA affects how they function. Models of this fold use two kinds of data: contact maps constructed from Hi-C, and nucleosome maps showing where DNA is wound around nucleosomes. Over evolution, sections of the genome were copied, creating near-identical regions. Reads from these regions cannot be assigned to one copy and are discarded, leaving “void regions” in both maps. We tested MaaD-TAD, a pipeline that models topologically associating domains (TADs) at the atomic level, on the PRAMEF/HNRNPCL locus on human chromosome 1. This challenging locus spans 652 kb, with only 15% of its nucleosome positions known and most contact data missing. To test our hypothesis that copies with similar sequence lie close in space, i.e. homology as a surrogate for proximity, we constructed a map of sequence homology in place of Hi-C and used CSynth to fold it into a 3D path. We then lengthened the path to fit a chromatin fibre, and refined the model by adding cohesin complexes, which stabilize chromatin structure. Cohesin is a ring-shaped protein complex through which a loop of chromatin is threaded; the loop is commonly anchored by CTCF proteins bound on either leg. Using known databases, we placed CTCF sites in our model and joined three to their copies with cohesin rings; in each pair the sites face each other, as loop anchors do. The final model holds 3,359 nucleosomes and about 55 million atoms. Micro-C, an advanced, higher-resolution form of Hi-C, provided data on two other duplicated regions that support our hypothesis: matching copies touched 1.2 to 1.9 times more often than expected. Our approach provides a novel method to reconstruct DNA regions with absent contact data and, to our knowledge, the first all-atom model of a complete TAD.

**Author summary:** Inside every cell, two metres of DNA are folded into a nucleus a few thousandths of a millimetre across, and where a gene sits in this fold helps decide whether it is used. We map the fold by reading millions of short DNA fragments, but regions of the genome that were copied during evolution look the same to these methods, so their data are thrown away. We asked whether such a blank region could still be modelled. We focused on a region of human chromosome 1, 652,000 DNA letters long, that holds two families of copied genes, PRAMEF and HNRNPCL, which are switched on in the earliest stages of the human embryo. Our working idea was that copies with similar sequence lie close together in space, so we used sequence similarity in place of the missing data. We then built MaaD-TAD, a pipeline that builds a 3D model in which every atom of the DNA and every atom of its packaging proteins has a position, about 55 million atoms in all. The model shows how copied genes and their control switches could pair up, and makes predictions that experiments can test. The same pipeline also builds models from measured maps.

## Introduction

### Why build an all-atom model of a chromatin domain

Each human cell packs roughly two metres of DNA into a nucleus a few micrometres across. To fit, the DNA wraps around protein spools called nucleosomes, forming a fibre known as chromatin, which is then folded in three dimensions. This fold is not random, and it may help regulate which genes are switched on. Each gene has a promoter, a stretch of DNA at and just upstream of where transcription begins. Genes are often switched on when an enhancer, a regulatory stretch of DNA, touches the promoter through a loop in the chromatin fibre. The enhancer can lie tens to hundreds of kilobases (kb) away. These contacts occur largely within topologically associating domains (TADs), neighbourhoods of the genome where DNA sequences contact one another more often than they contact DNA outside the domain. TAD borders are set by two proteins: CCCTC-binding factor (CTCF), which binds a short DNA sequence, and cohesin, a ring-shaped protein complex that can hold two strands of DNA together [1–3].

Most of what we know about this fold comes from chromosome conformation capture (Hi-C). In Hi-C, DNA that touches inside the cell is chemically joined, cut, and sequenced. Sequencing produces reads: short stretches of sequence, about 50 to 150 base pairs long, read from each end of a joined fragment. The result is a contact map, a table that reports how often each pair of regions meets. A contact map tells us that two regions meet, but not how. It does not say which nucleosomes are involved, which stretch of linker DNA (the free DNA between two nucleosomes) bridges them, or which face of the fibre they use. Hi-C maps are also coarse. To count contacts, the genome is divided into equal pieces called bins, and contacts are tallied bin by bin, much as a census tallies households by postal code. The bin size is chosen during analysis. Smaller bins show more detail but collect fewer contacts each, so their size depends on how many reads were sequenced. Typical bins are a few kb, a span of 10 to 25 nucleosomes.

Micro-C, a newer form of Hi-C, cuts DNA with the enzyme micrococcal nuclease, which trims the linker DNA between nucleosomes [4]. Each joined fragment is therefore about one nucleosome long, which sets the finest detail Micro-C can reach, although Micro-C maps are still usually counted in bins of a few kb. Even so, every map is an average over millions of cells, a probabilistic consensus that smooths out differences between individual cells.

Three sizes recur in this paper, and it helps to tell them apart. A read (about 100 bp) is a short piece of sequence. Its length decides whether it can be placed in the genome, not how much detail a map shows. A nucleosome (about 190 bp, counting its linker) is the physical unit that our model places. A bin is the counting unit chosen for a map. In this study bins are 2 kb, about 10 nucleosomes, for every map of the locus, matching the sequence-similarity (SEQSIM) map described below [5]. In the census analogy, a read is one ballot, a nucleosome is one house, and a bin is the postal code used for the tally.

An all-atom model of a TAD, in which the position of every atom is known, would let us ask how two regions meet, not only whether they do. Can an enhancer and its promoter physically touch without their atoms overlapping or their molecular bonds being pulled apart? Can two specific DNA segments pass through a single cohesin ring given their physical proximity? Here we describe MaaD-TAD (Molecular all-atom Display of TADs), a pipeline that builds such a model for an entire TAD. It starts from a contact map and a nucleosome map and ends with every base pair and every histone protein placed at atomic detail. After its first step, the pipeline works on a three-dimensional path rather than on the map itself. It can therefore start with three kinds of input. The first is a measured Hi-C or Micro-C map (Fig 10). The second is the SEQSIM map of sequence similarity, for regions where measured data are missing. The third is a hybrid, in which similarity fills the gaps of a measured map. To our knowledge, the result is the first all-atom model of a complete TAD.

### The data gap

Most TADs cannot yet be modelled at the atomic level, because the data are incomplete. Hi-C and nucleosome maps are both built from short sequencing reads. Each read must be matched to one unique place in the reference genome, the standard human DNA sequence. Some regions of the genome, however, are segmental duplications: long stretches that were copied, so that two or more near-identical versions exist. A read from such a region matches several places equally well. It cannot be assigned, so it is discarded from further analysis. One can picture a jigsaw puzzle in which several pieces show exactly the same patch of sky. Each piece fits more than one spot, so it is set aside. The result is a blank stripe in the contact map (Fig 1B) and a hole in the nucleosome map (Fig 1A).

The same regions are often poorly assembled. The region studied here, the PRAMEF/HNRNPCL locus (see below), still contains a gap of 50 kb in the current human reference genome (GRCh38). Its length also changed by 40% between versions of the same reference, from 913 kb in the previous version (GRCh37) to 652 kb in the current version, GRCh38. Yet duplicated regions are exactly where gene families expand, where the number of copies of a gene varies between people, and where rearrangements of the chromosome recur [6, 7]. They are among the TADs most in need of a structural description, and the least served by current data.

### Sequence similarity as a stand-in for contact

Where contact data are missing, we propose to use sequence homology as a proxy for spatial proximity. Two stretches of DNA are homologous when they share an ancestor, which shows as a similar sequence. Although we are not aware of a study that uses homology this way, several independent lines of evidence make it plausible. We reason from three premises and two supporting observations.

#### Premise 1: copies arise between neighbours

When DNA is copied from one place to another, the donor and recipient sites are usually close to each other. When the machinery that copies DNA before cell division stalls, it can switch to a nearby template and copy it instead [8]. Pieces of chromosomes that swap places in disease, called translocation partners, tend to be regions that already touch in the nucleus [9]. Viruses that insert their DNA into ours follow the same geography: HIV-1 inserts preferentially near nuclear speckles, bodies in the nucleus rich in gene activity [10]. A pair of homologous copies is therefore a record that the two places were once close to each other. Homology can be read as a fossil of past proximity. This is clearest in clusters formed by repeated tandem duplication, where new gene copies start side by side. The question then becomes whether they stay close as new copies push them apart.

#### Premise 2: copied anchors hold the pair together

When a DNA segment is duplicated, any CTCF binding site within it is copied as well. Bursts of mobile DNA elements, called retrotransposons, have spread CTCF sites through mammalian genomes and reshaped their loops [11]. Direction matters. The CTCF binding sequence, or motif, reads in one direction, like an arrow. When a copy is inverted, the arrow flips, so the copy and its source either face each other (a convergent pair) or point away from each other (a divergent pair). Loops held by cohesin form mainly between convergent sites [1–3]. Copying alone can therefore create the anchor for a loop, without any new binding site.

#### Premise 3: being close is useful

Genes that arose by duplication within one genome are called paralogs. Paralogs are found in the same TAD more often than expected. Paralogs more than 1,000 kb apart also touch each other more often than pairs of unrelated genes at the same distance [12]. They share enhancers more often (27% of pairs, against 12% for unrelated genes) and are often switched on together [12]. Promoters held side by side could draw on the same transcription factors, the proteins that turn genes on, within a small, shared space inside the TAD. This benefit would favour keeping the copies close once they arise, so that they are regulated together.

### Supporting observations: DNA can recognize matching DNA

Intact double-stranded DNA pairs with matching double-stranded DNA, even when no proteins are present [13]. The attraction is weak for each base pair but grows with the length of the match [14]. The copies studied here are 97.9–99.98% identical over tens of kb. At the scale of the whole nucleus, repeats of the same family cluster together and help divide the genome into compartments, large active and inactive zones [15].

Together, these observations suggest that homologous regions are more likely than random pairs to lie close together. This would be true both because of how the copies arose and because their closeness can be maintained. We treat this as a working hypothesis. It provides the contact map that the data cannot, and it makes predictions that can be tested.

### The PRAMEF region as a test case

To test the pipeline under the hardest conditions, we chose the PRAMEF/HNRNPCL locus, a region of 652 kb on the short arm of human chromosome 1 (band 1p36; GRCh38 chr1:12,771,346–13,423,345). A locus is simply a defined position on a chromosome. This one is named after its two gene families: PRAMEF (PRAME family members, where PRAME stands for preferentially expressed antigen in melanoma) and HNRNPCL (heterogeneous nuclear ribonucleoprotein C-like). The PRAME family expanded independently in several lineages of placental mammals [16]. In humans, the locus holds 26 protein-coding genes (22 PRAMEF and 4 HNRNPCL) and 8 pseudogenes, broken gene copies that no longer make protein. These are arranged in sixteen homology blocks, pairs of copied stretches that are 97.9–99.98% identical (Fig 1C). In twelve blocks the copy is inverted, and in four it runs in the same direction, a direct copy. Only 15% of the nucleosomes in the locus have measured positions, and the locus is blank in public Hi-C maps (Fig 1A and 1B).

Little is known about what these genes do, but what is known points to early development and to disease. PRAME, the founding member of the family, is a tumour antigen that represses retinoic acid receptor signalling [17] and acts as the substrate-recognition subunit of a Cullin-2 ubiquitin ligase [18]. PRAMEF genes are among the genes that the transcription factor DUX4 switches on at the cleavage stage of the human embryo, when the embryo’s own genome first becomes active [19]. DUX4 is not expressed in healthy muscle, but it is expressed in the muscle of people with facioscapulohumeral muscular dystrophy (FSHD), where it activates PRAMEF1; in muscle cells made to express DUX4, it also activates HNRNPCL1 [20]. In mice, loss of Pramef12 arrests sperm production and makes males sterile [21]. The HNRNPCL genes, named for their similarity to the RNA-binding protein hnRNP C, are otherwise largely uncharacterized. The locus is therefore a cluster of copied genes that one transcription factor switches on together, which makes the arrangement of its copies in space a biological question as well as a technical one. We replaced the missing contact map with a sequence-similarity map, made with our previously developed SEQSIM approach [5]. SEQSIM scores how alike the sequences of every pair of bins are, on both strands, and lays the scores out in the same grid as a Hi-C contact map. We folded this map into a three-dimensional path with CSynth [22]. CSynth is an interactive program that turns a contact map into a chain of beads, one bead per bin, like beads on a string (Fig 2). We then show that this path, as drawn, cannot hold a real chromatin fibre: it is too short to hold the nucleosomes of the locus even as a compact fibre, parts of it pass closer to each other than the fibre is wide, and it bends more tightly than the fibre can. We refolded it to account for the thickness and stiffness of the fibre (a step we call Folder; Fig 3). Nucleosomes were then placed along the refolded path as a two-start zigzag, the arrangement in which consecutive nucleosomes alternate between two stacks (Placer). Every linker DNA was then built at atomic detail and joined to the next nucleosome (Builder), and each section of the model was checked and repaired (Checker; Figs 4 and 5). Finally, we held homologous arms of the model parallel and in register (Fig 6). Two copies are in register when each position on one copy faces the matching position on the other. Picture a step ladder: the two copies are the side rails, and each rung joins a matching pair. In register, every rung is level and the same length. Out of register, one rail has slid along the other, so the rungs slant and join positions that do not match. For an inverted copy, one rail simply runs in the opposite direction. We also joined three pairs of copied CTCF sites with cohesin rings (Fig 7). The result is an all-atom model of 652,000 base pairs of DNA on 3,359 nucleosomes. The standard method for mapping where a protein binds DNA is chromatin immunoprecipitation sequencing (ChIP-seq). The same homology search also predicts fifteen CTCF sites that ChIP-seq missed, because they lie in sequence where reads cannot be placed (Table 1).

S1 Table lists the key terms used in this paper, with a short plain-language explanation of each.

## Materials and methods

### Genomic region, sequence and gene annotation

The PRAMEF/HNRNPCL locus was defined as GRCh38 chr1:12,771,346–13,423,345 (652,000 bp). Its sequence was retrieved from the UCSC Genome Browser REST interface (hg38) and contains a 50 kb run of N at 12,954,385–13,004,384, an unresolved assembly gap. The same genes span chr1:12.83–13.75 Mb (913 kb) in GRCh37 (Ensembl GRCh37 REST lookup).

Genes were taken from Ensembl (GRCh38, release 116) through its REST interface, using the canonical transcript of each gene. The locus contains 26 protein-coding genes (22 *PRAMEF*, 4 *HNRNPCL*) and 8 pseudogenes; gene coordinates matched NCBI RefSeq Select exactly. Sixteen homology blocks (97.9–99.98% identity; 12 inverted, 4 direct) were identified (manuscript in preparation). Each block consists of two near-identical copies, which we call its arms. In a direct block the second arm repeats the first in the same orientation, so the two arms can lie side by side only if the DNA between them forms a parallel loop, a loop whose arms run in the same direction. In an inverted block it is the reverse complement, so the two arms run in opposite directions and fold back on each other as a hairpin. We call the hairpin’s turning point its tip; in the sequence-similarity map it is where the stripe of an inverted block meets the diagonal (Fig 2C). Genes that occupy matching positions on the two arms of a block are called mirror-image pairs (Fig 6C). Blocks are labelled in genomic order I1–I12 (inverted) and D1–D4 (direct).

### Notation

The labels used throughout the paper are listed in Table 2.

### Nucleosome positions

The locus was assigned 3,376 nucleosomes. Measured positions of the nucleosome dyad (the central base pair of the DNA wrapped on the histone octamer) were available for 490 (15%), all within the first third of the locus. These positions came from MNase-seq of the lymphoblastoid cell line GM12878, taken from the GM12878 nucleosome signal of ENCODE/Stanford/BYU in the UCSC Genome Browser (track wgEncodeSydhNsomeGm12878Sig) [23]. The track exists only on hg19 (GRCh37), so its positions were converted to GRCh38 coordinates with liftOver and the UCSC hg19–hg38 chain. As a check, the 490 dyads were lifted back to hg19 (verify_nucleosome_source.py, S1 File): 80% lay within 10 bp of a local maximum of this track, against 9% of random positions, and their scores correlated with its signal (Spearman ρ = 0.94). Where alternative positions overlapped, one was kept so that no nucleosome repeat length (the distance between consecutive dyads) fell below 163 bp, for two reasons. First, 163 bp (the 147 bp core plus a 16 bp linker) is the shortest repeat that the two-start zigzag template used for placement can represent. Second, a dyad map is an average over many cells, so two dyads called less than a core apart more likely mark alternative positions of one nucleosome, each used in a different fraction of cells, than two neighbours on the same fibre. This affected 130 of 3,420 linkers; in each case the stronger peak (higher occupancy) was kept, and a measured dyad was always kept over a placeholder. The remaining positions were filled with placeholders at a repeat length of 190 bp, within the range measured in primary human cells (178–203 bp, depending on cell type and chromatin domain) [24]. Together with the measured positions, this gives a mean repeat of 194 bp over the locus, the spacing used in Fig 3. Of all nucleosomes, 2,468 (73%) lay in sequence that could not be mapped uniquely. Opening naked DNA at the three cohesin tethers removed 17 nucleosomes, leaving 3,359 in the final model.

### SEQSIM contact map

In place of a Hi-C map, we obtained pairwise sequence-similarity scores using our previously developed SEQSIM approach [5] (alignment-based similarity between each pair of sections). A 1.3 Mb window centred on the locus was divided into 650 sections of 2 kb; each pair of sections received a similarity score that included the reverse complement, so that inverted copies were detected. The 26 sections overlapping the assembly gap were omitted. Sections 163–488 (chr1:12,771,321–13,423,320) were used to fold the locus (Fig 2A).

As an independent check of the SEQSIM scores, we also scored each pair of bins by the percentage of 90 bp words they share on either strand. This simpler score agreed with SEQSIM (Spearman correlation 0.93, log scale, over homologous bin pairs) and recovered every pair with a SEQSIM score above 20.

### Three-dimensional path

The SEQSIM map was folded into a path of 326 beads, one per 2 kb section, with CSynth [22] (latest web version, csynth.org; accessed September 2026) using its default settings. CSynth places a bead in every 2 kb bin of the folded range, including bins without data, and exports each bead with its bin position. Beads 91–116, which fall in the assembly gap, had no similarity data, so only CSynth’s backbone and non-overlap terms acted on them. They form a smooth arc between the two flanking beads, with gentler turns than the rest of the path (4–21° between successive steps, against a median of 15° elsewhere). Their positions are therefore a geometric placeholder, not a prediction.

### MaaD-TAD

All steps after CSynth were carried out with MaaD-TAD (Molecular all-atom Display of TADs), a Python pipeline written for this study (S1 File). It has four modules, run in order (Fig 4A). **Folder** refolds the CSynth path so that it can hold a chromatin fibre, and applies the homology pairing restraints and tether terms (folder.py). **Placer** places the nucleosomes along the refolded path (placer.py). **Builder** builds the DNA and histones in atoms and closes every linker (builder.py). **Checker** runs the structural checks and rebuilds the blocks that fail them (checker.py). Each module is described below.

### Folder: refolding the path for a chromatin fibre

A CSynth path has no physical units. Before refolding, it was scaled so that its two most distant beads were 500 nm apart. This span comes from fluorescence in situ hybridization (FISH) imaging of the locus (manuscript in preparation). It is an adjustable parameter of Folder, so users can set it from their own imaging data.

A path drawn from a contact map has no thickness, and a chromatin fibre folded on it would pass through itself (Fig 3). The path was therefore refolded by energy minimization (autograd gradients and the limited-memory Broyden–Fletcher–Goldfarb–Shanno (L-BFGS) minimizer of SciPy [25]) with four constraints:

- Excluded volume: no two fibre segments closer than 52 nm.
- Stiffness: bend radius of at least 130 nm (90 nm allowed within 6 kb of a tether), a safety margin well above the fibre’s physical minimum of about 48 nm (Fig 3B).
- Length: the arc length of a spline through the path matched the fibre length required by the nucleosomes it carries, within a window of 1.000–1.005.
- Fidelity: a restraint toward the CSynth path (weight 0.02), acting only on bead pairs closer than 300 nm on the CSynth path.

Homology pairing and tether terms were added at this stage (see below). From this step on the path has physical units: its largest span was 963.5 nm after refolding and 1,011.5 nm once the pairing and tether terms were added. All parameters of Folder, Placer, Builder and Checker are listed in S2 Table.

### Placer: nucleosome placement

Nucleosomes were placed along the refolded path as a two-start zigzag fibre (radius about 11.3 nm, 138.7° twist per nucleosome step), with the geometry of each step taken from a nine-nucleosome two-start zigzag built with G-Dash (Genome Dashboard, version available in 2025) [26], the longest fibre the software exports. They were added one at a time, each obtained from the previous one by the rotation and translation of one zigzag step, steered to follow the path.

Before each placement was accepted, the next four steps were tested, and placements were rejected if they over-stretched a linker (extension above 0.93 of its contour) or brought two nucleosomes within 12 nm.

Because each 2 kb section of the path was given the length its nucleosomes need, this step-by-step placement stayed close to the genomic position of each nucleosome (median offset 7 nm; 95% within 23 nm). Each nucleosome was also anchored to its genomic position on the path, as a guard against slip, which kept each nucleosome a median of 11 nm from its genomic position along the path (95% within 20 nm; S1 Fig). Registration was measured as the arc-length offset between the point of the path nearest each nucleosome centre and the path point for its dyad, with genomic position mapped linearly between the 2 kb beads. The median steering rotation applied per nucleosome was 2.8° (95th percentile 9.5°; S1 Fig).

### Builder: all-atom construction

Each nucleosome core was built from the crystal structure of the nucleosome core particle (PDB 1KX5) [27], 147 bp of DNA wrapped on a histone octamer. Linker DNA between consecutive cores was built one base-pair step at a time. Step parameters were expanded in smooth bending and twisting modes and solved by damped Gauss–Newton iteration so that the linker closed onto the next core, to within 10⁻¹⁰ nm. The solver started with several seeds (arcs and S-curves of varied curvature and direction) and ranked the closed solutions first by clearance from nearby cores, treated as discs, and then by strain. Failed linkers were retried with more bending modes and seeds. The sugar–phosphate backbone of each linker strand was then built from a library of 1KX5 conformations and joined to the flanking cores. Finally, the GRCh38 sequence of the locus was threaded onto the model (thread_sequence.py, S1 File), because the cores and linkers are built from template bases. Chain A residue r corresponds to genomic position r + 12,771,345 and carries the reference base; chain B residue r corresponds to position 13,423,346 − r and carries its complement. Each base that differed from the reference was replaced by superposing a base of the new type on the old one by C1′ and three atoms of the glycosidic ring (purine N9, C4 and C8; pyrimidine N1, C2 and C6). The sugar–phosphate backbone, the linker closures and the block seams are therefore unchanged. Bases inside the 50 kb assembly gap (N in GRCh38) kept their template identity. The real sequence on the model is needed for any later test of base pairing between homologous copies. Threading changed 900,222 of the 1,204,000 bases outside the gap (75%). Afterwards all 652,000 base pairs were Watson–Crick pairs, and the median distance between the paired N1 and N3 atoms was 0.29 nm, as in B-DNA. Because a new base is sometimes larger than the one it replaces, atom pairs closer than 0.22 nm between a base and other DNA rose from 50,038 to 67,953 (compare_base_contacts.py, S1 File). Close contacts between bases and histones were already present before threading (335,972 before, 332,654 after) and were removed in the next step.

### Histone tails

The histone tails are disordered in cells, but 1KX5 resolves each in a single conformation, and copying that conformation onto every core placed the N-terminal tail of one H3 copy (chain E, residues 1–39) across the linker DNA entering the nucleosome, where the crystal has no DNA. Every tail with an atom within 0.22 nm of DNA was therefore moved by rotating only its backbone φ and ψ angles, from the folded domain outward (repair_tails.py, S1 File). Each angle was set on a 10° grid to the value that kept the tail farthest from both DNA and other protein atoms (up to 0.36 nm and 0.32 nm), with three sweeps per tail and a second, finer pass (5°, six sweeps) for the three tails that still touched DNA. Rotations about backbone bonds leave every bond length and bond angle unchanged (largest change 10⁻⁶ nm), and the histone folds and the DNA did not move. The repair moved 6,380 tails, 3,291 of them the H3 tail of chain E, and reduced tail atoms within 0.22 nm of DNA from 230,918 to 4, all on one H3 tail.

Afterwards no base lay within 0.22 nm of a histone atom, and no tail lay within 0.22 nm of another protein. Because each block was repaired on its own, three H3 tails had been swung into chromatin belonging to another block; these were relaxed again with the neighbouring blocks held fixed (aiming for 0.45 nm from DNA and 0.40 nm from protein; repair_tails.py --from-qc), after which the whole-model check below found no long-range clashes. The 67,953 remaining atom pairs closer than 0.22 nm between a base and other DNA lie between neighbouring nucleotides, a base and the backbone of the next nucleotide in a bent linker or a base and its pairing partner; this is local strain of the kind that energy minimization removes (see Limitations).

The model, about 55 million atoms, was built in 68 blocks of 50 nucleosomes in parallel (on a single Windows workstation with an Intel Core i9-14900K processor (24 cores) and 128 GB of RAM), and blocks were joined at seams by shared linkers. The block size is a practical choice, not a structural one: each linker is solved from the positions of its two cores and of the cores around it, whichever block it falls in. Fifty nucleosomes (about 10 kb of DNA and 0.8 million atoms, a 72 MB mmCIF file) is small enough to build in one process and to open and inspect on its own in PyMOL. The resulting 68 blocks keep 24 parallel processes busy, and a repair rebuilds only the blocks whose nucleosomes moved. Block size is an option of Builder (S1 File). Residues were numbered by genomic position: chain A residue *r* = genomic position − 12,771,345; chain B residue *r* = 13,423,346 − genomic position.

### Folder: homology pairing restraints and cohesin tethers

#### Pairing

For each of the 16 homology blocks, the two arms were restrained during refolding on three counts. Their spacing was held near its value on the CSynth path. They were kept parallel, for direct repeats, or antiparallel, for inverted repeats. And they were kept in register: lines joining homologous positions were held perpendicular to both arms (weights 10 for direction and register, and 2 for spacing).

#### Register error

To measure how well each homology block was paired, each path was resampled at 20 points per 2 kb bead by linear interpolation. For every fifth point along the first copy, we found the nearest point on the second copy. The register error of that point was the genomic distance between the nearest point and the matching position, which runs in the same direction along the second copy for direct repeats and in the opposite direction for inverted repeats. The median over each copy gave one value per block. Three paths were compared: the CSynth path, the path refolded without pairing restraints, and the final path carrying pairing restraints and tethers (Fig 6B). The cost of pairing to the overall fold was measured as the Spearman correlation of pairwise bead distances between each refolded path and the CSynth path.

#### Tethers

Three cohesin tethers (T_I2, T_I5, T_I7, named after the inverted block in which each lies) each joined a peak of the cohesin subunit RAD21 to a mirror copy of the same CTCF site with reported CTCF binding (Table 1). At each site the DNA was left free of nucleosomes over at least 400 bp, centred on the site. Both duplexes must pass through the same ring, so each needs a stretch of naked DNA long enough to reach the ring and leave it again with its flanking nucleosome cores (11 nm across) kept outside; 400 bp gives about 68 nm of naked DNA on either side of the site, more than the 20 nm radius of the ring used during building. Because removing a nucleosome lengthens the linker between its neighbours, the naked stretches in the final model were 422–746 bp. CTCF-bound sites also lie in nucleosome-depleted regions in cells [28]. Within a tether pinch of 6 kb, the pairing terms were relaxed so that the arms converged only at the tether (local pinch).

A cohesin ring was modelled as a torus of radius 20 nm and tube radius 1.2 nm, through which the two naked duplexes passed with their axes 3.5 nm apart. Each tether linker was solved in two halves that met at a waypoint on the ring. The half built second avoided the partner’s finished backbone, with its axis at least 1.9 nm from any partner phosphorus, and the histone Cα atoms of nearby cores, with its axis at least 1.4 nm from any Cα. The ring drawn in Fig 7C was fitted to the finished model (fig7c.pml, S1 File): a torus of tube radius 1.0 nm was moved and tilted to maximize its clearance from all atoms, with the condition that the two naked duplexes, and no other atoms, pass through its opening. The smallest radius meeting this condition was found by testing radii from 4.0 nm upwards. For T_I5 the ring radius was 4.5 nm (opening 7.0 nm), the duplex axes passed 1.6 and 1.8 nm from its centre and 3.45 nm from each other, and the closest atom was 0.97 nm from the ring surface; for T_I7 and T_I2 the radii were 4.5 and 5.5 nm, the axis separations 3.29 and 3.62 nm, and the clearances 0.93 and 0.89 nm.

### Checker: structural quality control and repair

The finished model was checked for completeness of both strands; covalent continuity, meaning that every O3′–P bond was present, including those across block seams; phosphodiester geometry, counting O3′–P–O5′ angles outside 95–115° as strained; and long-range steric clashes between parts more than five nucleosomes apart (P–P < 0.5 nm, Cα–P < 0.4 nm, Cα–Cα < 0.38 nm), found with k-d trees, a data structure for fast searches of nearby atoms.

Clashes were repaired iteratively. A coarse stand-in of the model was built: DNA axes, with a minimum of 2.2 nm between two axes and 1.1 nm between an axis and a histone Cα. Flagged nucleosomes were moved apart on this stand-in by rigid-body relaxation, the ring waypoints were recomputed, the stand-in was re-checked against the new waypoints, and only blocks whose frames had changed were rebuilt. In the final model both strands were complete (652,000 residues each). All 1,303,998 O3′–P bonds were intact, 0.01% of linkages were strained, and no long-range clashes remained.

### Cohesin and CTCF sites and prediction of mirror sites

ChIP-seq peaks for cohesin subunits (RAD21, SMC3, SMC1A, STAG1, STAG2) and CTCF were collected from three sources through the UCSC Genome Browser REST interface:

Encyclopedia of DNA Elements (ENCODE) transcription factor ChIP-seq clusters (track encRegTfbsClustered) [29], ENCODE candidate cis-regulatory elements flagged as CTCF-bound (encodeCcreCombined), and ReMap 2022, a database of published ChIP-seq peaks (ReMapTFs) [30]. Nine sites had cohesin peaks and ten had CTCF peaks without cohesin (Table 1).

The best match to the CTCF core motif was found within each peak with the CTCF matrix MA0139.2 from JASPAR, a database of transcription-factor binding profiles (JASPAR 2026 [31], read with pyJASPAR and Biopython 1.88 [32]). The matrix records how often each base occurs at each position of known CTCF sites. Every stretch of the peak was scored, on both strands, by how much better than chance its bases fit the matrix (a log-odds score; a pseudocount of 0.5 was added to every count so that a base never seen at a position does not score minus infinity). The best score in each peak was then rescaled from 0 (the worst possible match) to 1 (a perfect match), as 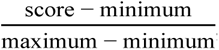, and reported with its strand. Homologous copies of each site were found by aligning ±150 bp around its best motif to the whole locus with minimap2 [33] (mappy 2.31, short-read preset), keeping copies with at least 90% identity over at least 80% of the window. A copy with no ChIP-seq peak within 300 bp was classed as a predicted mirror site: the copy of a known CTCF site, which ChIP-seq cannot test because reads there cannot be placed. This gave 15 predicted mirror sites: 6 copies of cohesin sites and 9 copies of CTCF-only sites. Motif orientation at pairs of ChIP-supported copies was classed as convergent or divergent (Fig 7D).

### Hi-C analysis of control loci

#### Choice of control loci

To test whether homologous sequences lie closer together in space, we selected duplicated loci in which paralogous copies have diverged enough for sequencing reads to be placed uniquely. Four candidate windows were retrieved from the UCSC Genome Browser (GRCh38) through its REST interface: the protocadherin cluster (chr5:140,700,001– 141,600,000), a zinc-finger gene cluster (chr19:57,000,001–58,000,000), the Th2 cytokine locus containing *IL13* (chr5:132,300,001–133,000,000) and the β-globin locus with its flanking olfactory receptor genes (chr11:5,000,001–5,700,000). The *IL13* and β-globin windows contained 0 and 4 homologous bin pairs and were not analyzed further.

#### Homology maps

Each window was divided into 2 kb bins. Interspersed repeats, soft-masked in the UCSC sequence, were removed, and bins with fewer than 300 unmasked bases were excluded. Every pair of bins was aligned on both strands by Smith–Waterman local alignment (parasail 1.3.4 [34]; match +1, mismatch −3, gap opening 5, gap extension 2). The higher of the two scores was divided by the length of the shorter unmasked bin and expressed as a percentage. Bin pairs scoring 30 or more were considered homologous: 133 pairs in the protocadherin cluster and 41 in the zinc-finger cluster. For folding, scores were rescaled to the SEQSIM scale by quantile matching against the PRAMEF SEQSIM map, on which a score of 30 corresponds to a SEQSIM value of 1.1. To compare how alike the copies are, each homologous pair was re-aligned with a traceback, and its identity was taken as the share of matching bases over the aligned columns, gaps included; the PRAMEF window was scored the same way (S3 Table; control_identity.py, S1 File).

#### Micro-C data

We used Micro-C data from H1 human embryonic stem cells [4], obtained from the 4D Nucleome Data Portal [35] (experiment set 4DNES21D8SP8; contact matrix 4DNFI9GMP2J8; GRCh38). Balanced contact frequencies in 2 kb bins (about 10 nucleosomes each; the same bin size as the homology maps) were read directly from the multi-resolution cooler file with h5py 3.16.0. The file was produced with a cooler version that can list a pixel more than once; for such pixels the largest count was kept. Contacts were converted to observed/expected (O/E) values by dividing each entry by the mean of its diagonal.

#### Mappability of the locus (**Fig 1B**)

The same Micro-C file was used to show how contact maps behave over the locus. Raw counts and balancing weights in 2 kb bins (about 10 nucleosomes each; the bin size of the SEQSIM map) were extracted for chr1:11,800,000–14,400,000, the locus with about 1 Mb of flank on each side. Bins with an undefined balancing weight (discarded by the 4DN pipeline for low coverage) were counted, and read coverage per bin was taken as the row sum of raw counts within the window. Inside the locus, 227 of 327 bins (69%) were discarded, against 3 of 973 in the flanks, and median coverage was 37-fold lower than in the flanks (97 vs 3,544 contacts per bin). Fig 1B shows balanced contacts on a log scale, clipped at the 5ᵗʰ and 99.3ʳᵈ percentiles of non-zero values.

#### Contact enrichment test

For each homologous bin pair separated by at least three bins and covered by Micro-C, one control pair at the same genomic separation was drawn at random from pairs with a homology score below 9. This left 122 homologous pairs in the protocadherin cluster and 39 in the zinc-finger cluster. The O/E values of homologous and control pairs were compared with a one-sided Mann–Whitney U test (SciPy 1.17.1), and enrichment was reported as the median of the per-pair ratios (Fig 8B).

#### Three-dimensional paths

Homology-guided paths were built with a polymer model that follows the design of CSynth [22], re-implemented in Python. Beads of 2 kb were joined by springs of unit rest length (stiffness 50). Each pair of beads was attracted by a harmonic term proportional to its homology score above background, scaled by a homology weight *k*, and repelled by a soft term 0.3/(*d*² + 0.25), where *d* is the distance between beads. The energy was minimized by L-BFGS (SciPy) from random-walk starting conformations, and the lowest-energy result of four starts was kept. On the PRAMEF SEQSIM map, this model reproduced the CSynth path used in this study with a Spearman correlation of pairwise distances of 0.76–0.86, similar to the agreement between two independent runs of the model (0.85).

Micro-C contacts were folded with the same model after scaling balanced counts so that their 99.9ᵗʰ percentile equalled 100. Homology-guided and Micro-C-guided paths were compared by the Spearman correlation of pairwise distances between bins at least three apart, with folds built without homology restraints as a baseline. Contacts predicted from the homology-guided path (proportional to *d*⁻³) were compared with Micro-C O/E values by Spearman correlation.

#### Calibration of the homology weight (**Fig 8C**)

The homology map of each control locus (on the SEQSIM-like scale, homologous pairs score above 1.1 and control pairs at most 0.6) was folded with a CSynth-like model (csynth_like.py, S1 File): 2 kb beads joined by harmonic springs, a soft repulsion between non-adjacent beads, and a harmonic attraction between bin pairs weighted by their similarity above the map median and scaled by the homology weight *k*, minimized with L-BFGS from a random-walk start. Weights *k* = 0, 0.002, 0.005, 0.01, 0.02 and 0.05 were tested with five random starts each, with the attraction applied either to homologous pairs only or to the whole map; *k* = 0 is a chain with springs and repulsion only. For each path, contact enrichment was the median ratio of distance⁻³ between each homologous pair and a control pair at the same separation, and the calibrated weight was the *k* at which the mean enrichment met the Micro-C value of Fig 8B (linear interpolation). Agreement with the path folded from Micro-C (compare_hic.py --fold) was measured after removing the shared dependence on genomic separation: log distances were converted to z-scores within each separation, and the two paths were compared by Pearson correlation (sweep_calibrate.py, S1 File). The PRAMEF homology map was folded in the same way at *k* = 0 and *k* = 0.007, and the same enrichment was measured in the CSynth path, the refolded path and the path of the final model (pramef_calibration.py, S1 File).

#### Mapping test of the 145 kb line (**Fig 9**)

To estimate how many contacts on the line could come from reads assigned to the wrong copy, reads of 50, 100 and 150 bp were taken every 10 bp along each 5 kb bin of the line (13.225–13.245 Mb and 13.370–13.390 Mb), given 0.5% random substitutions as sequencing errors, and aligned to the GRCh38 sequence of the locus with minimap2 [33] (mappy 2.31, short-read preset). For each bin we counted the fraction of reads whose best alignment fell in the partner bin, 145 kb away, with mapping quality of at least 1 or at least 30 (misassign_pramef.py, S1 File). The expected number of misassigned pairs on the line was this fraction multiplied by the local contacts of the bin (raw counts of the bin with itself and its two neighbours), summed over both partners. Identity between partner bins was measured by aligning each 5 kb bin to the partner region (minimap2 asm20 preset). Reads were aligned to the locus only; copies elsewhere in the genome could remove reads from the locus but not add contacts to the line. To test whether the line is specific to H1 cells, the same bins were read from Micro-C of HFFc6 foreskin fibroblasts [4] (4D Nucleome Data Portal, 4DNFI643OYP9; GRCh38; 5 kb) with check_stripe.py (S1 File). For each bin on the line, the script reports the raw read pairs on the line, the mean of the four bins one step off it, the local contacts of each bin and its balancing weight. Read pairs expected from misassignment in HFFc6 were computed from the same simulated fractions (50 bp reads, mapping quality ≥1) and the HFFc6 local contacts.

For Fig 9B and C, extract_stripe_maps.py (S1 File) reads raw counts in 5 kb bins (used for the line instead of 2 kb so that each bin pair holds enough read pairs to compare with its neighbours) for chr1:13,100,000–13,500,000 from both files; the panels show rows 13.170–13.270 Mb against columns 13.315–13.415 Mb, coloured by log₁₀(count + 1) on the ramp used for Fig 1B, scaled to the maximum of each map. Fig 9A joins the 2 kb H1 map of Fig 1B (325 bins from 12,772,000, upper triangle, same colour limits) with the SEQSIM map of Fig 2A (lower triangle).

#### Validation of the analysis

The analysis was first tested on synthetic contact maps generated from known structures, with contact frequency proportional to *d*⁻³ and Poisson noise. Maps generated from the homology-guided path returned an enrichment of 5.1× (*P* = 6 × 10⁻³⁰) and a model–data correlation of 0.95. Maps generated from a fold without homology returned 1.0× (*P* = 0.67) and a correlation of −0.12.

### Builds from Micro-C alone (Fig 10)

The two test windows have near-complete Micro-C coverage and contain duplicated segments; they were taken from a scan of the H1 Micro-C map for lines of contacts parallel to the diagonal (manuscript in preparation). IGHC is chr14:105,540,001–105,750,000 (105 bins of 2 kb, 6 discarded by the 4DN pipeline) and PWRN is chr15:24,130,001–24,440,000 (155 bins, 7 discarded). Balanced H1 Micro-C counts in 2 kb bins were read from the file used for Fig 1B (extract_hic_test.py, S1 File), written as CSynth contact files with discarded bins left out (to_contacts.py, S1 File) and folded in CSynth [22]. SEQSIM maps of both windows were computed from the GRCh38 sequence in the same 2 kb bins (extract_hic_test.py, S1 File); they are shown for comparison in Fig 10A and were not used in the builds. Each path was then built with the PRAMEF settings (S2 Table) but without homology pairing or tethers (build_locus.py, S1 File), with two differences. First, CSynth units were scaled so that the mean bead spacing was 0.62 times the length of fibre supplied by the nucleosomes of one 2 kb bin, the same nucleosome density at the start of refolding as in PRAMEF. Second, the fidelity restraint acted on all bead pairs rather than only on pairs closer than 300 nm. These domains are 1.2–1.6 µm across, and with the 300 nm cut the 10 kb agreement of the full-atom models with H1 Micro-C fell to 0.47 (IGHC) and 0.25 (PWRN). Refolding PRAMEF with all pairs kept changed neither the pairing of its homology blocks nor its tethers (tethers 6.2–6.6 nm; every homologous arm pair still closer than the median at its separation), so the PRAMEF model was not rebuilt. No measured nucleosome positions were used for the test loci: nucleosomes were placed as a placeholder ladder with a repeat of 190–191 bp, chosen so that the chain spans each window exactly.

GRCh38 bases were threaded onto the blocks (thread_locus.py, S1 File) and the histone tails repaired as for PRAMEF. Agreement with Micro-C was measured as in the calibration above: log distances and log contacts were converted to z-scores within each genomic separation and compared by Pearson correlation, at 2 kb and on 10 kb blocks (5 × 5 bins, model contacts as mean distance⁻³) (score_models.py, S1 File). Contact enrichment of homologous bin pairs was measured as in the contact enrichment test above, with SEQSIM > 1.1 for homologous pairs and SEQSIM ≤ 0.6 for controls, in the H1 and HFFc6 Micro-C maps and in the model contacts of Fig 10B; the median ratio was stable across 200 random draws of the controls (homology_test_loci.py, S1 File).

### Figure preparation

Figure panels were drawn with Python scripts and rendered in PyMOL. The details needed to regenerate each panel, including input files, colour scales and camera settings, are given in S1 Text, and the scripts are in S1 File.

### Software, code and data availability

Analyses used Python 3.11 with NumPy 1.26.4, SciPy 1.17.1, autograd, gemmi, h5py 3.16.0 (3.11.0 for the Fig 1B extraction), Matplotlib 3.10.9, Pillow 12.2.0, parasail 1.3.4, mappy 2.31, Biopython 1.88 and pyJASPAR. Structures were inspected and rendered with PyMOL 3.1.4.1 (Schrödinger, LLC). The MaaD-TAD code, the Python pipeline that refolds the path and builds the all-atom model (Folder, Placer, Builder and Checker), is provided as S1 File and at https://github.com/mmarcet-palacios/MaaD-TAD (Zenodo, https://doi.org/10.5281/zenodo.23718187). The full-atom model, the nucleosome frames and the SEQSIM map are deposited in Zenodo (https://doi.org/10.5281/zenodo.23718429). Micro-C data are available from the 4D Nucleome Data Portal (H1, 4DNFI9GMP2J8; HFFc6, 4DNFI643OYP9).

## Results

### The PRAMEF/HNRNPCL locus lacks the data from which structural models usually start

The locus spans 652 kb of chromosome 1 (GRCh38 chr1:12,771,346–13,423,345) and carries 34 annotated PRAMEF and HNRNPCL genes and pseudogenes on both strands (Fig 1A). Given the nucleosome spacing used in the model, it holds 3,376 nucleosome positions. Only 490 of these (15%) have a measured dyad, and all 490 lie in the first third of the locus. A further 2,468 nucleosome positions (73%), covering 469 kb or 72% of the sequence, fall in sequence where short reads cannot be mapped uniquely (bracket). The locus also contains a 50 kb assembly gap (chr1:12,954,385–13,004,384; arrow).

Contact data fail over the same region (Fig 1B). Each bin of a Micro-C map collects the read pairs that touch it. Before the map is drawn, a correction step called balancing evens out bins that were sequenced more or less deeply. Bins with too few read pairs to correct are discarded and show as blank rows and columns. In the H1 Micro-C map, the flanks of the locus give a continuous map. Inside the locus, however, 227 of 327 bins (69%) were discarded, against only 3 of 973 in the flanks. The read counts show why. A typical (median) bin inside the locus holds 97 read pairs, against 3,544 in the flanks, 37 times fewer. Reads from the duplicated copies cannot be placed in the genome, so they never reach their bins. The blank stretch (12.864–13.364 Mb) overlaps the unmappable run marked in Fig 1A (12.954–13.423 Mb).

What the locus does have is extensive sequence homology (Fig 1C). Sixteen homology blocks join pairs of arms that are 97.9–99.98% identical: twelve are inverted repeats and four are direct repeats. ChIP-seq peaks from ENCODE and ReMap mark nine sites bound by cohesin and CTCF (C1–C9) and ten bound by CTCF alone (K1–K10). Searching the locus for copies of these sites found fifteen copies that keep an intact CTCF motif but have no reported peak: six copies of cohesin sites and nine of CTCF-only sites. Fourteen of the fifteen lie in sequence where short reads cannot be mapped uniquely. Table 1 lists the nine cohesin sites (C1–C9) with their homologous copies and the six predicted copies of cohesin sites (P1–P6). Six of the nine cohesin sites have at least one copy in the locus (C2, C3, C5, C6, C7 and C8), and all six predicted copies lie in unmappable sequence; P6 is the least similar to its source (94.1%).

**Fig 1.**
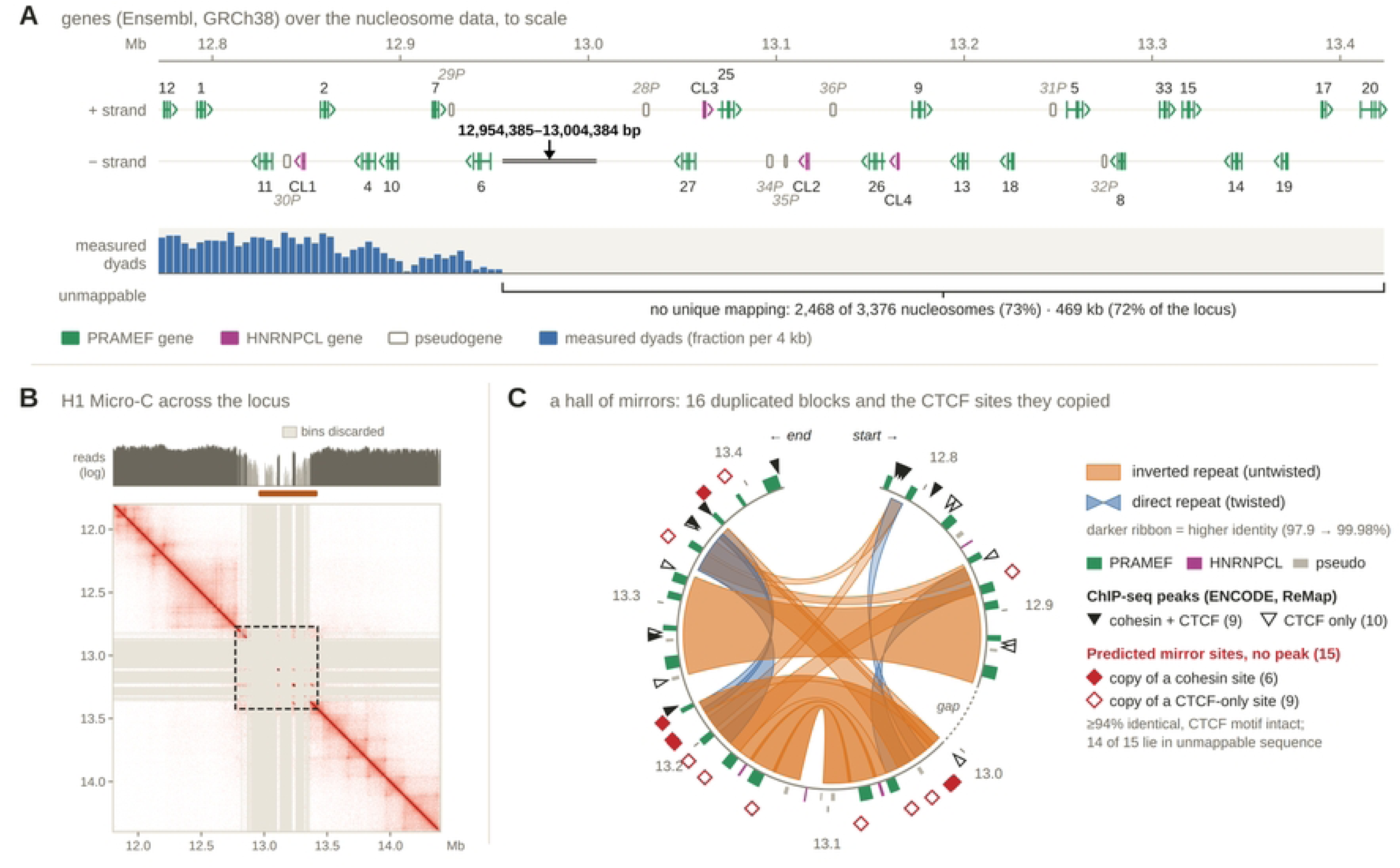
Gene annotation, nucleosome data, Micro-C contacts and sequence homology of the PRAMEF/HNRNPCL locus (GRCh38 chr1:12.77–13.42 Mb). (A) Protein-coding genes (Ensembl canonical transcripts, GRCh38; exons as boxes, chevrons point 5′→3′) and pseudogenes (outlined, italic labels) on the + strand (upper row) and − strand (lower row), drawn to scale above the nucleosome data; green, PRAMEF genes; magenta, HNRNPCL genes. Genes are labelled by number; in genomic order: PRAMEF12 (12), PRAMEF1 (1), PRAMEF11 (11), *PRAMEF30P* (30P), HNRNPCL1 (CL1), PRAMEF2 (2), PRAMEF4 (4), PRAMEF10 (10), PRAMEF7 (7), *PRAMEF29P* (29P), PRAMEF6 (6), *PRAMEF28P* (28P), PRAMEF27 (27), HNRNPCL3 (CL3), PRAMEF25 (25), *PRAMEF34P* (34P), *PRAMEF35P* (35P), HNRNPCL2 (CL2), *PRAMEF36P* (36P), PRAMEF26 (26), HNRNPCL4 (CL4), PRAMEF9 (9), PRAMEF13 (13), PRAMEF18 (18), *PRAMEF31P* (31P), PRAMEF5 (5), *PRAMEF32P* (32P), PRAMEF8 (8), PRAMEF33 (33), PRAMEF15 (15), PRAMEF14 (14), PRAMEF19 (19), PRAMEF17 (17), PRAMEF20 (20). Blue bars, fraction of nucleosome positions with a measured dyad in each 4 kb bin. Bracket, sequence that cannot be mapped uniquely with short reads (Methods), with the number of nucleosome positions and length it covers. Arrow, 50 kb assembly gap (N) in GRCh38, with its coordinates. (B) Micro-C contact map of H1 human embryonic stem cells [4] (4DN 4DNFI9GMP2J8; GRCh38; 2 kb bins, the bin size of the SEQSIM map in Fig 2, balanced with the weights supplied in the file; logarithmic colour scale) over chr1:11.8–14.4 Mb. Top track, reads per bin before balancing (logarithmic scale). Rust strip, the sequence without unique mapping (bracket in A). Grey rows and columns, bins discarded during balancing for too few reads. Dashed box, the locus. (C) Circular map of the locus, read clockwise from start to end. Ribbons join the two arms of each of the sixteen homology blocks: orange, untwisted ribbons, inverted repeats; blue, twisted ribbons, direct repeats; ribbon shade, identity between the two arms (darker, more identical; scale 97.9–99.98%). Outer ring, genes and pseudogenes as in A; dashed segment, the assembly gap (chr1:12,954,385–13,004,384). Black triangles, ChIP-seq peaks from ENCODE TF clusters [29] and ReMap 2022 [30]: filled, cohesin (RAD21 or SMC3) with CTCF; open, CTCF without cohesin. Red diamonds, predicted mirror sites: copies of a ChIP-supported site that retain the CTCF motif (JASPAR MA0139.2) but have no reported peak, found by aligning ±150 bp around each site to the locus (minimap2; ≥90% identity over ≥80% of the window); filled, copies of cohesin sites; open, copies of CTCF-only sites. The number of sites in each class is given in the key.

**Table 1.**
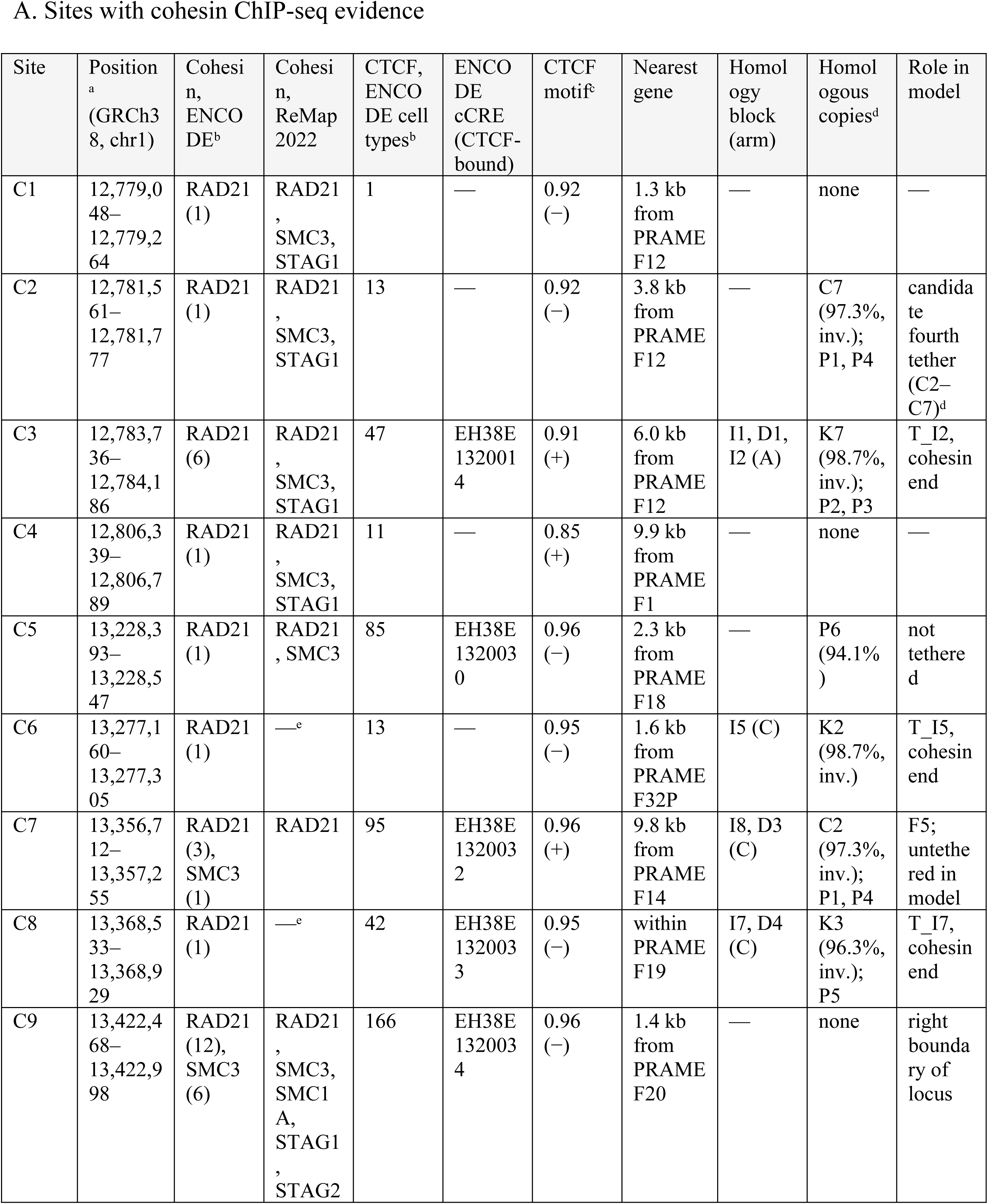
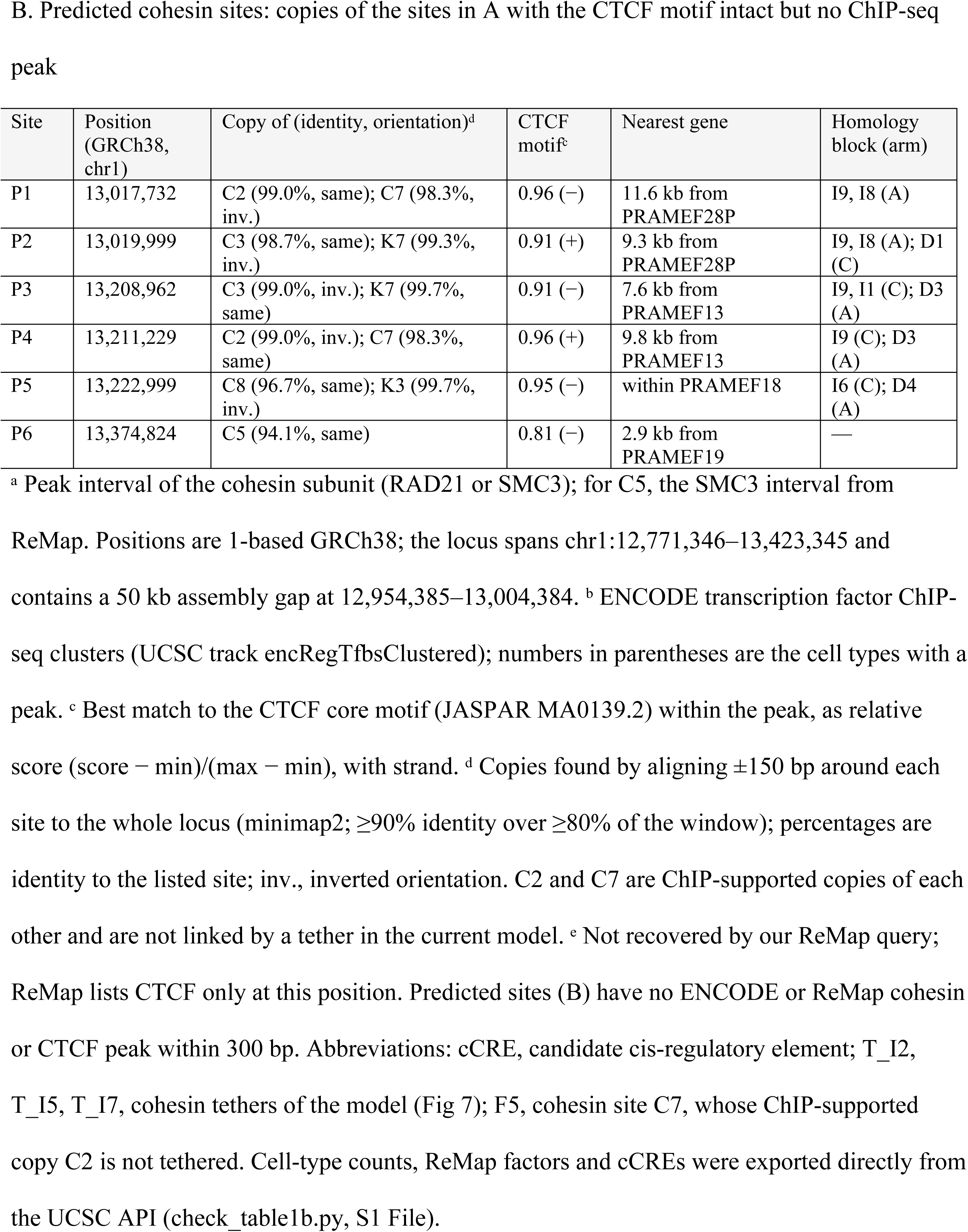
Cohesin-binding sites in the PRAMEF/HNRNPCL locus and their unannotated homologous copies.

| Site | Position<sup>a</sup> (GRCh38, chr1) | Cohesin, ENCODE<sup>b</sup> | Cohesin, ReMap 2022 | CTCF, ENCODE cell types<sup>b</sup> | ENCODE cCRE (CTCF-bound) | CTCF motif<sup>c</sup> | Nearest gene | Homology block (arm) | Homologous copies<sup>d</sup> | Role in model |
|---|---|---|---|---|---|---|---|---|---|---|
| A. Sites with cohesin ChIP-seq evidence |  |  |  |  |  |  |  |  |  |  |
| C1 | 12,779,048–12,779,264 | RAD21 (1) | RAD21, SMC3, STAG1 | 1 | — | 0.92 (−) | 1.3 kb from PRAMEF12 | — | none | — |
| C2 | 12,781,561–12,781,777 | RAD21 (1) | RAD21, SMC3, STAG1 | 13 | — | 0.92 (−) | 3.8 kb from PRAMEF12 | — | C7 (97.3%, inv.); P1, P4 | candidate fourth tether (C2–C7)<sup>d</sup> |
| C3 | 12,783,736–12,784,186 | RAD21 (6) | RAD21, SMC3, STAG1 | 47 | EH38E1320014 | 0.91 (+) | 6.0 kb from PRAMEF12 | I1, D1, I2 (A) | K7 (98.7%, inv.); P2, P3 | T_I2, cohesin end |
| C4 | 12,806,339–12,806,789 | RAD21 (1) | RAD21, SMC3, STAG1 | 11 | — | 0.85 (+) | 9.9 kb from PRAMEF1 | — | none | — |
| C5 | 13,228,393–13,228,547 | RAD21 (1) | RAD21, SMC3 | 85 | EH38E1320030 | 0.96 (−) | 2.3 kb from PRAMEF18 | — | P6 (94.1%) | not tethered |
| C6 | 13,277,160–13,277,305 | RAD21 (1) | —<sup>e</sup> | 13 | — | 0.95 (−) | 1.6 kb from PRAMEF32P | I5 (C) | K2 (98.7%, inv.) | T_I5, cohesin end |
| C7 | 13,356,712–13,357,255 | RAD21 (3), SMC3 (1) | RAD21 | 95 | EH38E1320032 | 0.96 (+) | 9.8 kb from PRAMEF14 | I8, D3 (C) | C2 (97.3%, inv.); P1, P4 | F5; untethered in model |
| C8 | 13,368,533–13,368,929 | RAD21 (1) | —<sup>e</sup> | 42 | EH38E1320033 | 0.95 (−) | within PRAMEF19 | I7, D4 (C) | K3 (96.3%, inv.); P5 | T_I7, cohesin end |
| C9 | 13,422,468–13,422,998 | RAD21 (12), SMC3 (6) | RAD21, SMC3, SMC1A, STAG1, STAG2 | 166 | EH38E1320034 | 0.96 (−) | 1.4 kb from PRAMEF20 | — | none | right boundary of locus |
| B. Predicted cohesin sites: copies of the sites in A with the CTCF motif intact but no ChIP-seq peak |  |  |  |  |  |  |  |  |  |  |
| Site | Position (GRCh38, chr1) | Copy of (identity, orientation)<sup>d</sup> | CTCF motif<sup>c</sup> | Nearest gene | Homology block (arm) |  |  |  |  |  |
| P1 | 13,017,732 | C2 (99.0%, same); C7 (98.3%, inv.) | 0.96 (−) | 11.6 kb from PRAMEF28P | I9, I8 (A) |  |  |  |  |  |
| P2 | 13,019,999 | C3 (98.7%, same); K7 (99.3%, inv.) | 0.91 (+) | 9.3 kb from PRAMEF28P | I9, I8 (A); D1 (C) |  |  |  |  |  |
| P3 | 13,208,962 | C3 (99.0%, inv.); K7 (99.7%, same) | 0.91 (−) | 7.6 kb from PRAMEF13 | I9, I1 (C); D3 (A) |  |  |  |  |  |
| P4 | 13,211,229 | C2 (99.0%, inv.); C7 (98.3%, same) | 0.96 (+) | 9.8 kb from PRAMEF13 | I9 (C); D3 (A) |  |  |  |  |  |
| P5 | 13,222,999 | C8 (96.7%, same); K3 (99.7%, inv.) | 0.95 (−) | within PRAMEF18 | I6 (C); D4 (A) |  |  |  |  |  |
| P6 | 13,374,824 | C5 (94.1%, same) | 0.81 (−) | 2.9 kb from PRAMEF19 | — |  |  |  |  |  |

<sup>a</sup> Peak interval of the cohesin subunit (RAD21 or SMC3); for C5, the SMC3 interval from ReMap. Positions are 1-based GRCh38; the locus spans chr1:12,771,346–13,423,345 and contains a 50 kb assembly gap at 12,954,385–13,004,384. <sup>b</sup> ENCODE transcription factor ChIP-seq clusters (UCSC track encRegTfbsClustered); numbers in parentheses are the cell types with a peak. <sup>c</sup> Best match to the CTCF core motif (JASPAR MA0139.2) within the peak, as relative score (score − min)/(max − min), with strand. <sup>d</sup> Copies found by aligning ±150 bp around each site to the whole locus (minimap2; ≥90% identity over ≥80% of the window); percentages are identity to the listed site; inv., inverted orientation. C2 and C7 are ChIP-supported copies of each other and are not linked by a tether in the current model. <sup>e</sup> Not recovered by our ReMap query; ReMap lists CTCF only at this position. Predicted sites (B) have no ENCODE or ReMap cohesin or CTCF peak within 300 bp. Abbreviations: cCRE, candidate cis-regulatory element; T\_I2, T\_I5, T\_I7, cohesin tethers of the model (Fig 7); F5, cohesin site C7, whose ChIP-supported copy C2 is not tethered. Cell-type counts, ReMap factors and cCREs were exported directly from the UCSC API (check\_table1b.py, S1 File).

### Sequence homology gives a contact map with two kinds of stripes, one folding as a parallel loop and the other as a hairpin

Because Hi-C contacts are missing for most of the locus, we used sequence similarity between 2 kb bins (SEQSIM; Fig 2A) in place of measured contacts. The sixteen homology blocks appear as stripes off the diagonal. Because similarity includes the reverse complement, a direct copy gives a stripe parallel to the diagonal and an inverted copy a stripe perpendicular to it. The assembly gap appears as grey rows and columns.

Fig 2B follows one direct copy, block D3 (98.8% identity), whose arms lie at 13.17–13.23 Mb and 13.32–13.38 Mb. Its stripe runs parallel to the diagonal. Two copies running in the same direction can only lie side by side if the 90 kb between them forms a parallel loop, a loop whose two arms run side by side in the same direction (middle). In the path folded from the map by CSynth (right), the two arms of D3 do run side by side in the same direction, with the intervening fibre looped out.

Fig 2C follows one inverted copy, block I12 (99.99% identity; 13.08–13.15 Mb). Its stripe crosses the diagonal at 13.113 Mb. Two copies running in opposite directions lie side by side when the fibre folds back on itself as a hairpin, with the tip where the stripe meets the diagonal (middle). In the CSynth path (right), the two arms of I12 run antiparallel and turn at 13.113 Mb.

**Fig 2.**
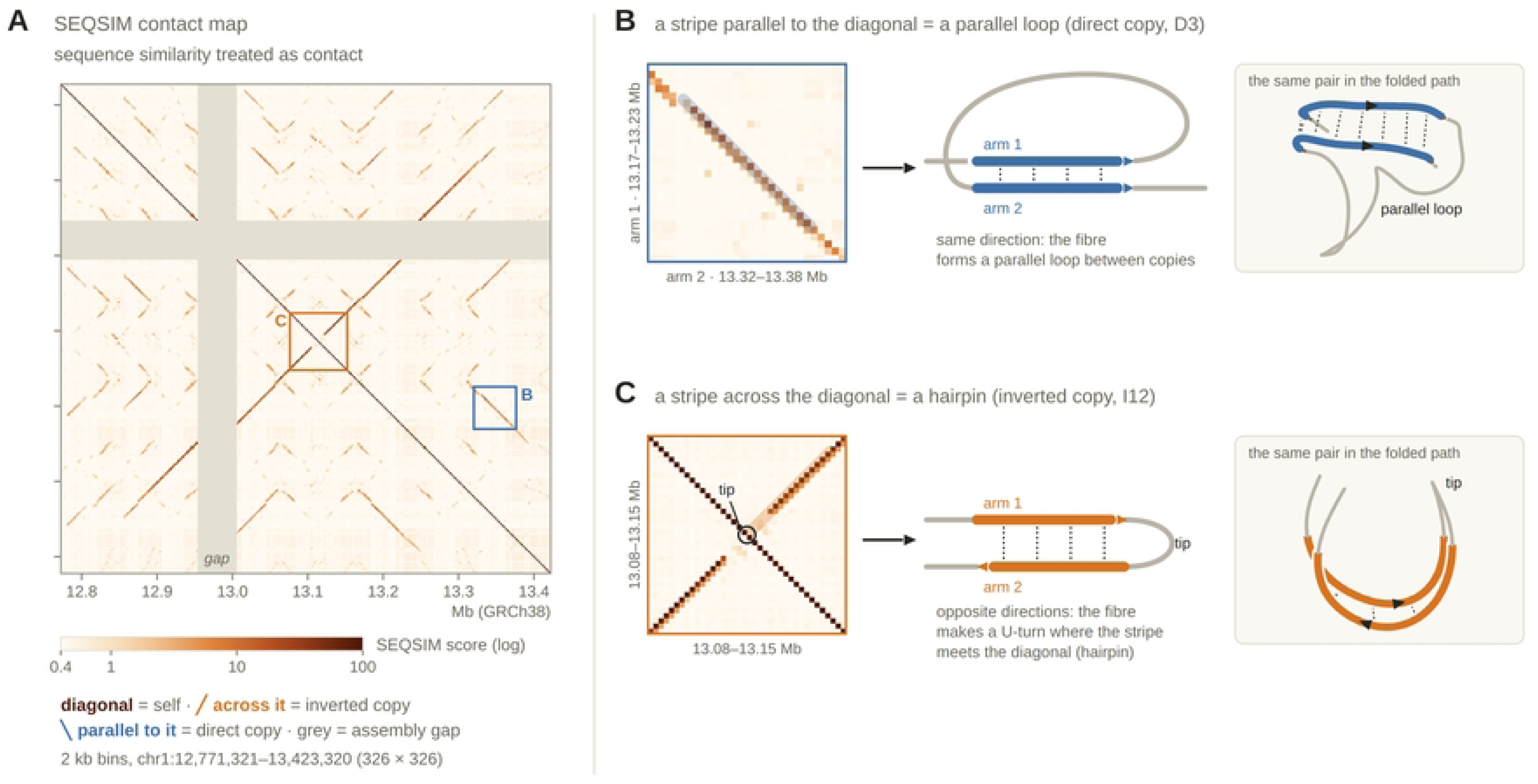
Homology-derived (SEQSIM) contact map of the locus, with one direct and one inverted homology block. (A) SEQSIM map of pairwise sequence similarity across the locus (chr1:12,771,321–13,423,320, GRCh38; 2 kb bins, 326 × 326; logarithmic colour scale, with colour key). Similarity includes the reverse complement. Grey rows and columns, 50 kb assembly gap (N). Boxes B and C, the regions enlarged in the left panels of B and C. (B) Block D3, a direct copy (98.8% identity). Left, the map enlarged on D3 (arm 1, chr1:13.17–13.23 Mb, rows; arm 2, 13.32–13.38 Mb, columns). Middle, schematic of two copies in the same orientation (arrows) joined by a parallel loop. Right, the same block in the path folded from the map by CSynth (2 kb beads), viewed slightly from above the plane of its two arms; dashed grey, fibre passing behind an arm; dotted lines join matching positions on the two arms; arrowheads, direction of each copy. (C) Block I12, an inverted copy (99.99% identity; chr1:13.08–13.15 Mb on both axes). Left, the map enlarged on I12. Middle, schematic of two copies in opposite orientations (arrows) joined at a tip. Right, the same block in the CSynth path, drawn as in B. Tip, the position where the stripe meets the diagonal (13.113 Mb).

### A chromatin fibre fits the homology-derived path only after the path is lengthened and refolded in 3D

The CSynth path gives the shape of the locus but not its length. Fig 3 compares the path with the fibre it must carry, in one, two and three dimensions. Scaled so that its two most distant points are 500 nm apart, the path is 9.1 µm long. Laid end to end at the spacing they have in the model (one every 23 nm; 194 bp repeat), the 3,359 nucleosomes would span 77.6 µm, 8.5 times the path (Fig 3A).

In a two-start zigzag, consecutive nucleosomes alternate between two stacks and the fibre axis advances 7.0 nm per nucleosome (Fig 3B). This brings the fibre down to 23.5 µm, still 2.6 times the path. The path is also too crowded for a 52 nm fibre: 113 of its 326 beads (35%) come closer than 52 nm to a non-adjacent part of the path (closest, 29 nm), and 18 bends have a radius below 48 nm.

We therefore refolded the path in 3D to the length of the fibre (Fig 3C). The refolded path is 23.4 µm long within a domain 0.96 µm across. No two parts come closer than 52 nm, the closest approach being between the two arms of the I12 hairpin (arrow), and no bend is tighter than 123 nm in radius (dashed circle). The refolded path keeps the CSynth fold (Spearman r = 0.91 between pairwise bead distances). Fig 3C shows the path before the pairing restraints and tethers described below (Figs 6 and 7); domain size and backfolding are analyzed in S2 and S3 Figs.

**Fig 3.**
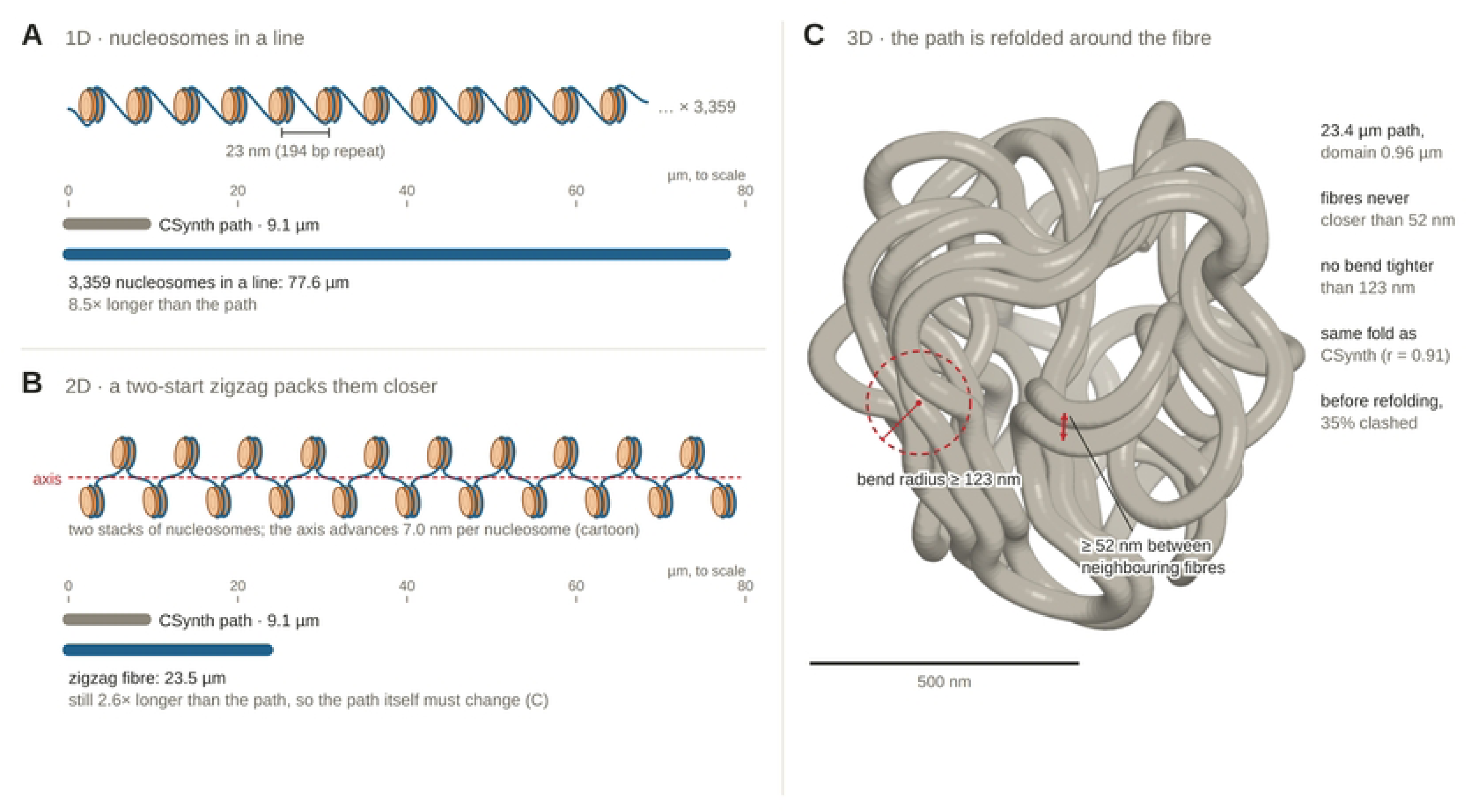
Length of the nucleosome chain and of the chromatin fibre compared with the CSynth path, and the path refolded in three dimensions. (A) Cartoon of nucleosomes (orange cylinders, the histone octamer; blue line, DNA making two turns around each octamer and continuing as linker DNA; colours as in Fig 5) laid in a line at the spacing used in the model (one every 23 nm; 194 bp repeat, bracket). Below, lengths drawn to scale on the µm axis: grey bar, the CSynth path, scaled so that its two most distant points are 500 nm apart (the span measured by FISH imaging of the locus; Methods); blue bar, the 3,359 nucleosomes of the locus laid in a line. (B) Cartoon of a two-start zigzag, in which consecutive nucleosomes alternate between two stacks; red dashed line, fibre axis. Below, lengths drawn to the same scale as in A: grey bar, the CSynth path; blue bar, the zigzag fibre with the axis advancing 7.0 nm per nucleosome. (C) The path refolded in 3D (Methods), shown before the pairing restraints and tethers of Figs 6 and 7 were added. Red double arrow, the closest approach between two non-adjacent parts of the path; red dashed circle and radius, one of the tightest bends. Scale bar, 500 nm. Right, values for the refolded path: length, domain diameter, minimum separation between non-adjacent parts, minimum bend radius, Spearman correlation of pairwise bead distances with the CSynth path, and the fraction of the CSynth path closer than 52 nm to a non-adjacent part before refolding. Domain-size and backfolding analyses are in S2 and S3 Figs.

### Five stages turn the homology map into a full-atom model

Fig 4A follows the locus through the pipeline, CSynth followed by the four modules of MaaD-TAD. The SEQSIM map (stage 1) is folded by CSynth into a 326-bead path (stage 2; each bead is a point standing for one 2 kb bin of the map, and consecutive beads are joined like beads on a string), which Folder refolds to carry a 52 nm fibre (stage 3; Fig 3). Placer then places the 3,359 nucleosomes along the refolded path as a two-start zigzag, each anchored to its genomic position (stage 4), and Builder builds the DNA and histones in atoms (stage 5), in 68 blocks of 50 nucleosomes. Checker then checks every bond and every long-range contact; flagged nucleosomes are relaxed and only the blocks that changed are rebuilt (return arrow). Placement needed a median steering rotation of 2.8° per nucleosome, and nucleosomes stayed a median of 11 nm from their genomic position along the path (95% within 20 nm; S1 Fig).

At the scale of a few nucleosomes, the model reproduces the zigzag it was built from. Fig 4B shows twelve consecutive nucleosomes (chr1:12,784,876–12,787,097) seen side-on to the fibre axis: consecutive nucleosomes alternate between two stacks on either side of the axis. Fig 4C enlarges one 40 bp linker. Each linker is built one base-pair step at a time from the exit of one core and lands on the entry of the next to within 10⁻¹⁰ nm, with backbone conformations taken from the nucleosome crystal structure (PDB 1KX5).

**Fig 4.**
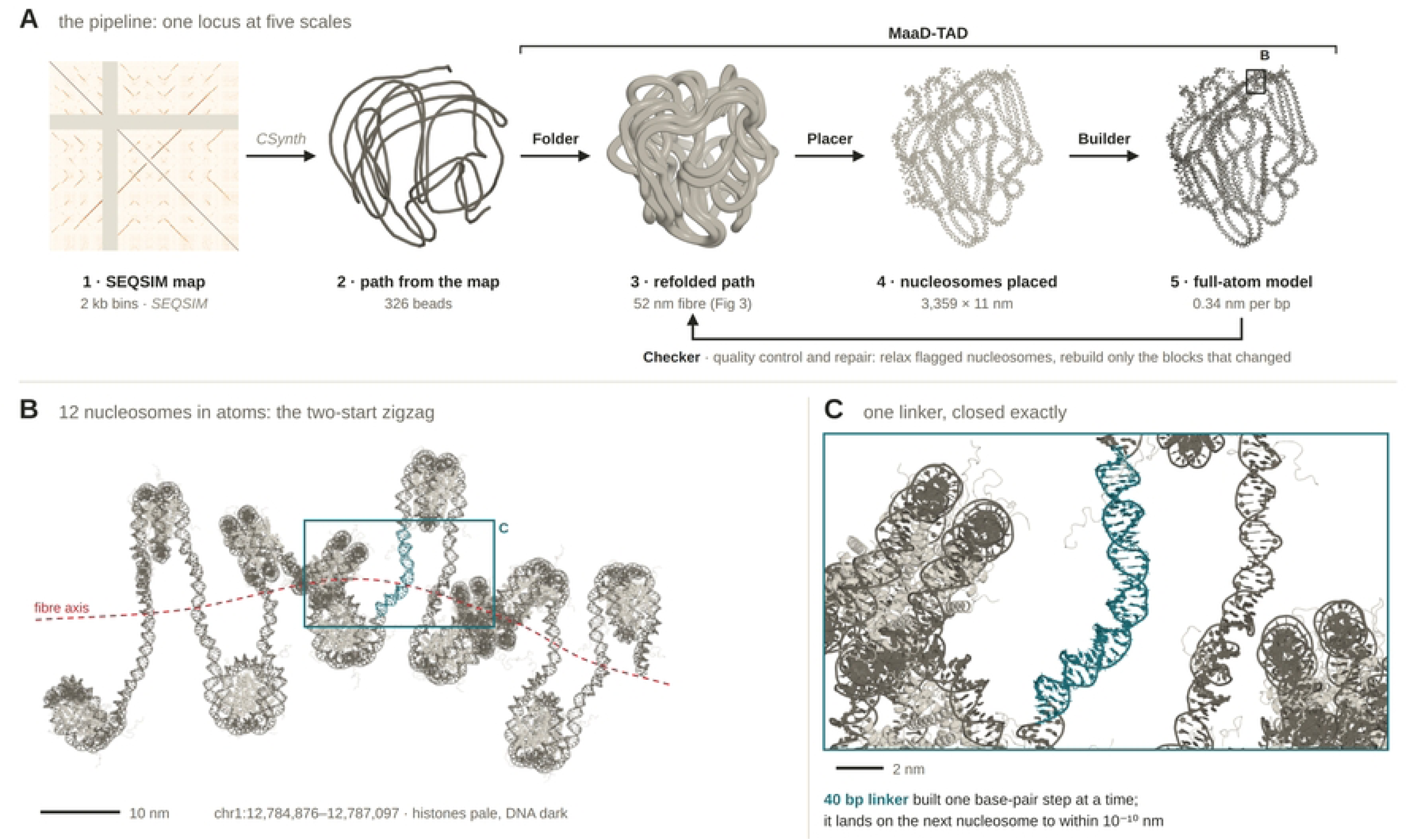
Pipeline from the homology map to the full-atom model, with two enlargements of the model. (A) The five stages, shown for the PRAMEF locus: (1) the SEQSIM map (2 kb bins; Fig 2); (2) the path folded from it by CSynth (326 beads); (3) the path refolded to carry a 52 nm fibre (Fig 3); (4) the 3,359 nucleosomes placed along the refolded path as a two-start zigzag, each anchored to its genomic position; (5) the full-atom model, built in 68 blocks of 50 nucleosomes (histones pale, DNA dark, as in B). Stages 2–5 are shown in one orientation. Labels above the arrows name the software for each step: grey italics, existing software (SEQSIM, CSynth); bold, the four modules of MaaD-TAD (Molecular all-atom Display of TADs; bracket): Folder, Placer, Builder and Checker (Methods; S1 File). Return arrow, the quality-control and repair loop run by Checker. Box, the region enlarged in B. (B) Twelve consecutive nucleosomes of the full-atom model (chr1:12,784,876–12,787,097), viewed side-on to the fibre axis (red dashed line; estimated from the nucleosome positions). Histones pale, DNA dark. Teal box, the region enlarged in C. Scale bar, 10 nm. (C) One 40 bp linker (teal) between two nucleosome cores. Scale bar, 2 nm.

### The final model packs 652 kb of chromatin into a domain about 1 µm across in which no DNA or histone passes through another part of the model

Fig 5 shows the final model at three magnifications along one viewing direction. The complete model (Fig 5A) holds 652,000 bp of DNA wrapped on 3,359 histone octamers, about 55 million atoms, and spans about 1 µm, the size of the refolded domain; S1 Video turns the complete model through 360°. Enlarged (Fig 5B), the domain resolves into neighbouring fibres lying side by side without interpenetrating, each a two-start zigzag. At the highest magnification (Fig 5C), single nucleosome cores and the linker DNA between them can be followed atom by atom.

The final build passed every structural check (Table 3). Both strands are complete (652,000 residues each), and all 1,303,998 O3′–P bonds are intact, including across the 67 seams between blocks. Only 0.01% of phosphodiester linkages are strained, there are no long-range clashes, no histone atom lies within 0.22 nm of a base, and every linker closes to within 10⁻¹⁰ nm. Nucleosomes lie a median of 11 nm from their genomic position along the path (95% within 20 nm; S1 Fig).

**Fig 5.**
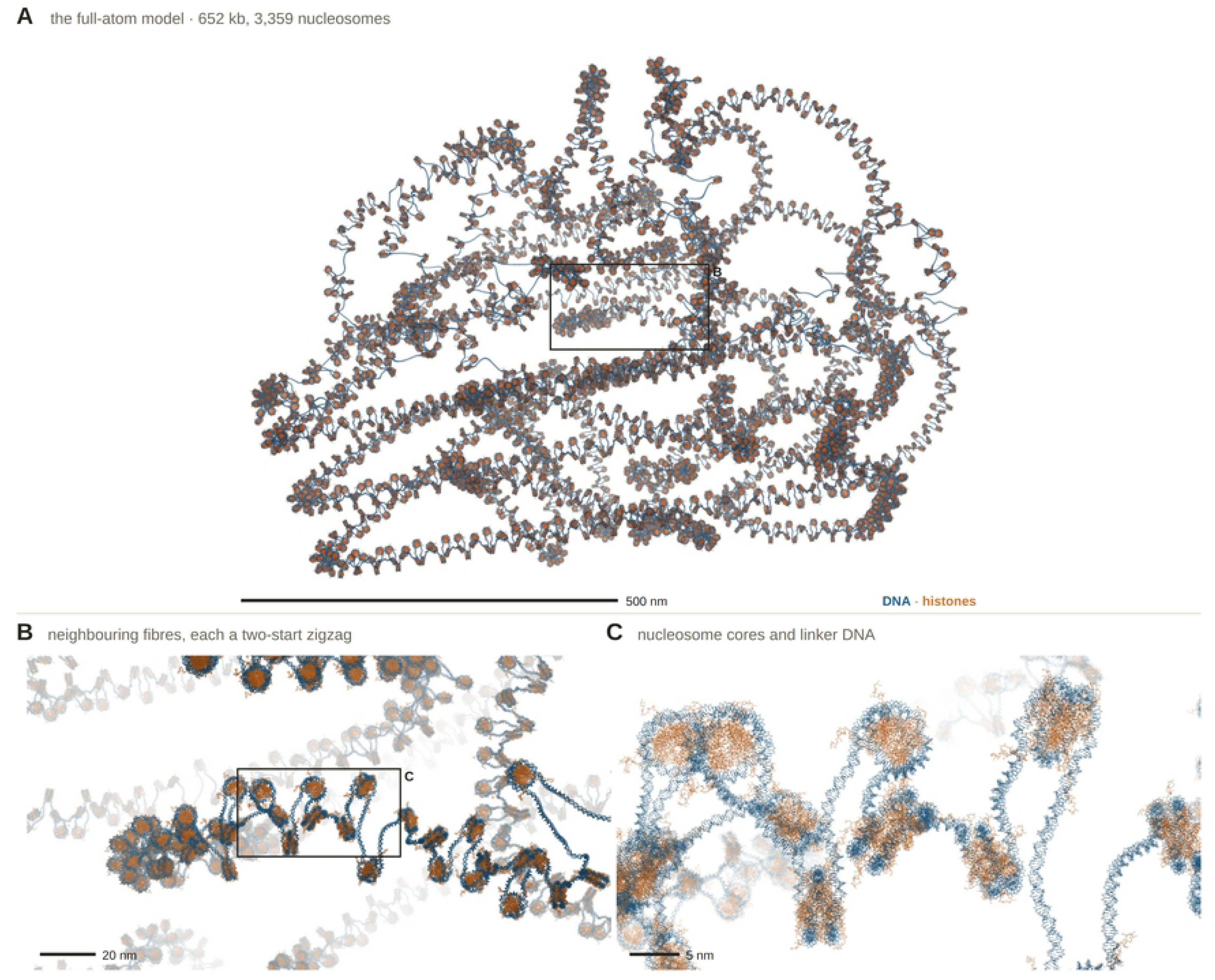
The full-atom model of chr1:12.77–13.42 Mb at three magnifications. (A) The complete model: DNA (blue) and histone octamers (orange), all atoms drawn as lines, rendered in PyMOL with orthographic projection and depth cueing on a white background. Box, the region enlarged in B. Scale bar, 500 nm. (B) Enlargement of the box in A. Box, the region enlarged in C. Scale bar, 20 nm. (C) Enlargement of the box in B. Scale bar, 5 nm. A–C share one viewing direction and centre; only the camera distance differs. Structural checks of the final build are given in Table 3.

**Table 2.** Labels used throughout the paper.

| Label | Meaning |
|---|---|
| I1–I12 | inverted homology blocks, numbered in genomic order |
| D1–D4 | direct homology blocks, numbered in genomic order |
| C1–C9 | sites with ChIP-seq peaks for cohesin and CTCF (Table 1A) |
| K1–K10 | sites with a ChIP-seq peak for CTCF only |
| P1–P6 | predicted copies of cohesin sites: intact CTCF motif, no ChIP-seq peak (Table 1B) |
| T_I2, T_I5, T_I7 | cohesin tethers of the model, each named after the inverted block that contains it |
| F5 | cohesin site C7, which has a ChIP-supported copy (C2) but no tether in the model |

**Table 3.** Structural quality of the final build. Checks are described in Materials and methods (Structural quality control and repair; Nucleosome placement).

| Check | Result |
|---|---|
| DNA residues per strand | 652,000 of 652,000 |
| O3′–P bonds intact | 1,303,998 (0 breaks) |
| Block seams joined | all 67, on both strands |
| Strained phosphodiester linkages (O3′–P–O5′ outside 95–115°) | 0.01% |
| Long-range clashes (P–P < 0.5 nm, Cα–P < 0.4 nm, Cα–Cα < 0.38 nm) | 0 |
| Base atoms within 0.22 nm of a histone atom (all atoms) | 0 (332,654 atom pairs before tail repair) |
| Base atoms within 0.22 nm of other DNA (all atoms) | 67,953 atom pairs, between neighbouring nucleotides (no force-field relaxation) |
| Linker closure error | < 10<sup>−10</sup> nm |
| Nucleosome registration (offset from genomic position along the path) | median 11 nm; 95% within 20 nm; maximum 55 nm, except the last 11 nucleosomes at the end of the chain (up to 140 nm; S1 Fig) |

### Homology pairing brings duplicated copies into register

The CSynth path places the two arms of each homology block near each other, but matching positions on the two copies are offset along the fibre. We measured this as a register error: for points along one copy, the distance along the partner copy between its nearest point and the matching position (Fig 6B). Across the 16 blocks, the median register error of the CSynth path was 2,018 bp, about ten nucleosomes (open circles). Refolding the path to fibre length did not change it (1,948 bp).

We therefore added a pairing restraint to the refolding (Fig 6A). Inverted copies are held antiparallel and direct copies parallel, and each rung joining two matching positions is kept at constant length and perpendicular to both copies. With pairing, the median register error fell to 133 bp, below one nucleosome repeat (194 bp; filled circles). The one block that stays out of register is D1, at 1.2 kb: its first arm is also an arm of I1 and I2, so one stretch of fibre has three partners. Pairing lowered the agreement with the CSynth fold from r = 0.91 to 0.80 and enlarged the domain slightly, from 0.96 to 1.01 µm (path length 23.2 µm).

Fig 6C shows what register means at the level of genes, in the nested I11/I12 hairpin (chr1:13.06–13.17 Mb) of the final model, viewed across the plane of the hairpin. Mirror-image genes on the two arms lie opposite each other: PRAMEF25 opposite PRAMEF26, the pseudogenes PRAMEF34P opposite PRAMEF36P, and HNRNPCL3 opposite HNRNPCL4. HNRNPCL2 sits at the tip, at the centre of the palindrome (13.113 Mb). Measured between nucleosome centres, the paired genes lie 83–109 nm apart centroid to centroid, and their closest nucleosomes 61–85 nm apart (PRAMEF25 and PRAMEF26, 88 and 64 nm; PRAMEF34P and PRAMEF36P, 109 and 85 nm; HNRNPCL3 and HNRNPCL4, 83 and 61 nm).

**Fig 6.**
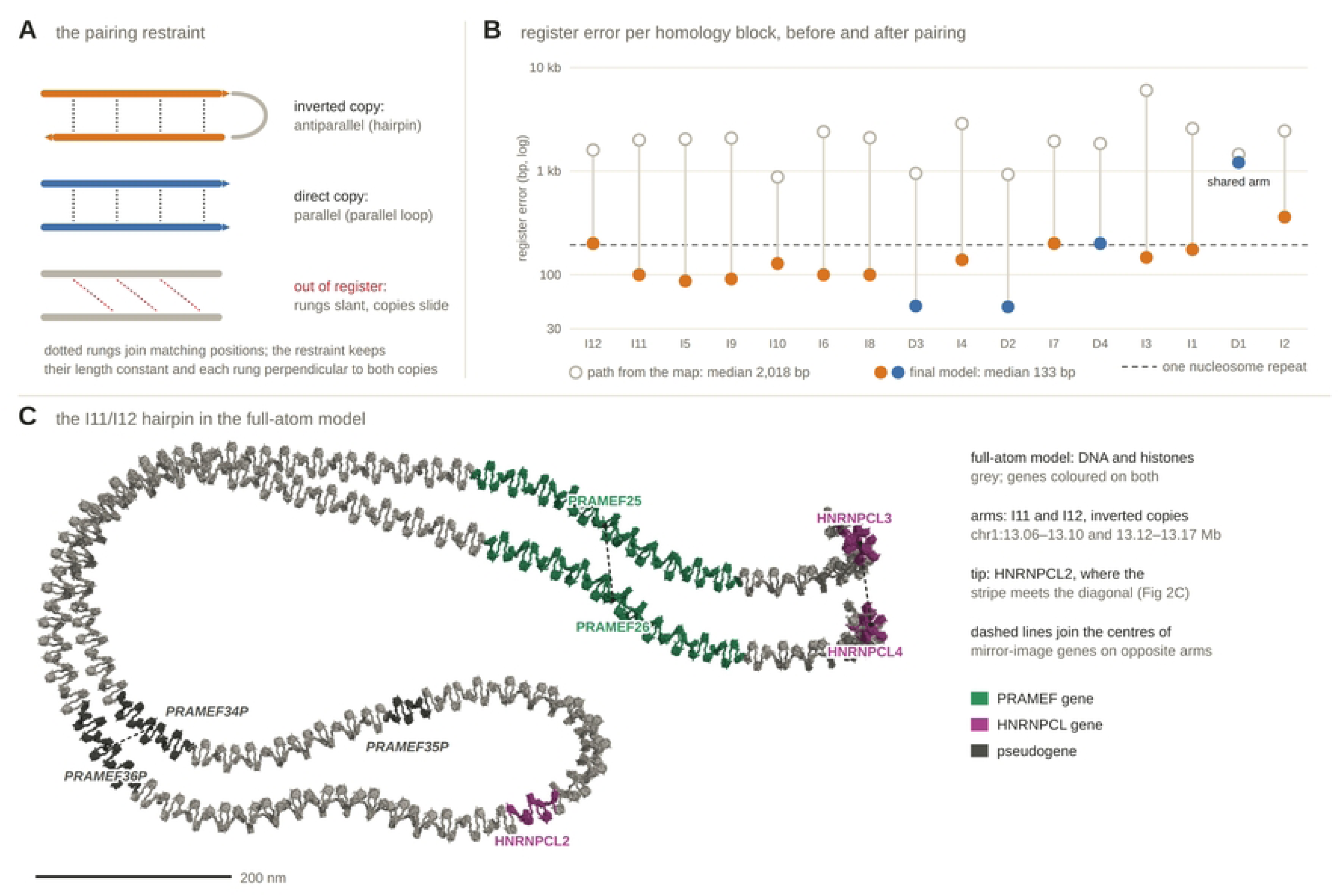
Homology pairing restraint, register error of the homology blocks, and the I11/I12 hairpin in the final model. (A) Schematic of the pairing restraint applied during refolding. Left, inverted copies held antiparallel (hairpin); middle, direct copies held parallel (parallel loop); right, two copies out of register. Dotted rungs join matching positions on the two copies; the restraint keeps each rung at constant length and perpendicular to both copies. (B) Register error of each of the 16 homology blocks (I, inverted; D, direct), defined as follows: for points along one copy, the distance along the partner copy between its nearest point and the matching position; median per block, logarithmic scale. Open circles, the path folded from the SEQSIM map by CSynth; filled circles, the path of the final model (orange, inverted copies; blue, direct copies); grey lines join the two values for each block. Dashed line, one nucleosome repeat (194 bp). Medians over the 16 blocks are given in the key. “Shared arm” marks D1, whose first copy is also an arm of I1 and I2. (C) The nested I11/I12 hairpin (chr1:13.06–13.17 Mb) in the full-atom model, viewed across the plane of the hairpin and rendered in PyMOL with orthographic projection on a white background. DNA and histones grey; genes coloured on their DNA and on the histones of nucleosomes whose dyad lies within them (green, PRAMEF; magenta, HNRNPCL; dark grey, pseudogenes). Dashed lines join the centres (dots) of mirror-image gene pairs on opposite arms; each centre is the centroid of the gene’s coloured region in the projection. The pseudogene PRAMEF35P has no mirror-image copy on the opposite arm. Scale bar, 200 nm.

### Three cohesin tethers join CTCF sites to their convergent homologous copies

Three cohesin sites have an inverted homologous copy that is itself bound by CTCF (Table 1), and the model tethers each site to its copy (Fig 7A). Sites are labelled as in Table 1 (C, sites with cohesin and CTCF peaks; K, sites with a CTCF peak only), and each tether is named after the inverted homology block that contains it (T_I2 lies in block I2; notation summarized in Methods). T_I2 joins C3 (12.784 Mb) to K7 (13.355 Mb), T_I5 joins C6 (13.277 Mb) to K2 (12.925 Mb), and T_I7 joins C8 (13.369 Mb) to K3 (13.006 Mb); the tethers span 571, 352 and 363 kb. The cohesin site C7 (F5, 13.357 Mb) has a ChIP-supported inverted copy, C2, but is not tethered.

Away from each tether the two arms keep their paired spacing, and they converge only at the tether itself (Fig 7B). At each tether site at least 400 bp of DNA is left free of nucleosomes, which removed 17 nucleosomes from the model (3,376 to 3,359). The tethers are the only places where fibres come closer than 48 nm: all nine such pairs of path points lie at tethers.

In atoms, both naked duplexes pass through a single ring, a schematic cohesin (Fig 7C). A ring of 4.5 nm radius (opening 7 nm across) is enough for T_I5 and T_I7, and one of 5.5 nm (opening 9 nm) for T_I2. Where they cross the plane of the ring, the two duplex axes are 3.3–3.6 nm apart (T_I5, 3.5 nm). In all three, no other DNA or histone passes through the opening, and the closest atom is 0.9–1.0 nm from the ring.

Five inverted pairs in the locus have ChIP-seq peaks at both copies (Fig 7D; Table 1). In the three tethered pairs the CTCF motifs are convergent (▶ ◀). In the two untethered pairs, C2–C7 and K1–K5, they are divergent (◀ ▶).

**Fig 7.**
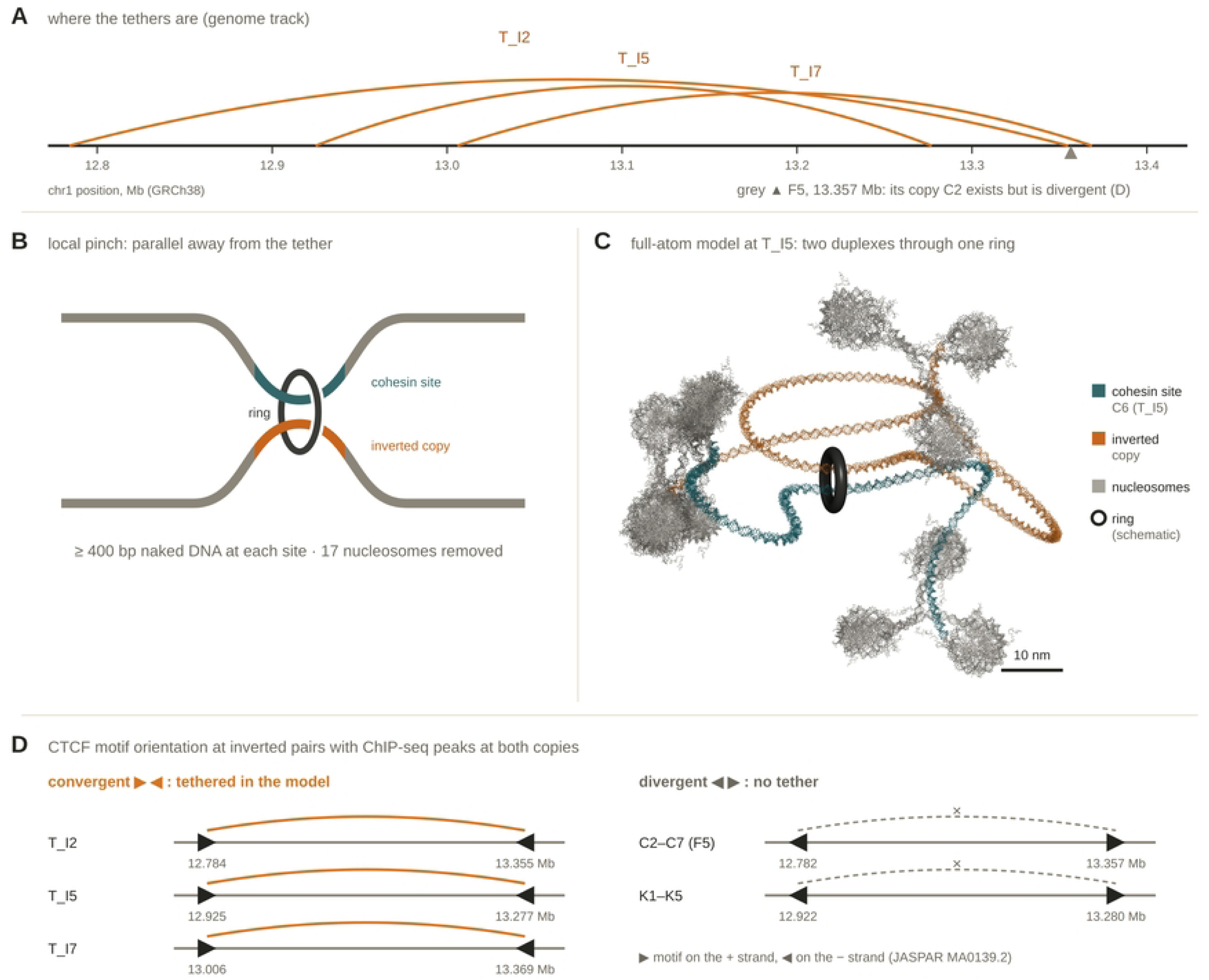
Cohesin tethers in the model and CTCF motif orientation at inverted pairs of sites. (A) Genome track of the locus (chr1:12.77–13.42 Mb, GRCh38; tick marks every 0.1 Mb) showing the three tethers of the model (orange arcs; T_I2, T_I5, T_I7), each joining a cohesin site to an inverted homologous copy with reported CTCF binding (Table 1). Grey triangle, cohesin site C7 (F5, 13.357 Mb), whose ChIP-supported copy C2 is not tethered. (B) Schematic of the fibre at a tether: two homologous arms (grey) pass through one cohesin ring (dark grey, as in C); teal, the cohesin site; orange, its inverted copy (colours as in C). (C) Full-atom model at tether T_I5 (PyMOL, all atoms as lines, ray-traced). Teal, DNA around cohesin site C6 (chr1:13,276,494–13,277,984); orange, DNA around its inverted copy (12,923,677–12,925,481); grey, the flanking nucleosomes (histones and DNA); black torus, a schematic cohesin ring (radius 4.5 nm, tube radius 1.0 nm) placed so that only the naked duplexes of the two sites pass through its opening (Methods). Scale bar, 10 nm. (D) CTCF motif orientation at the five inverted pairs of sites with ChIP-seq peaks at both copies (Table 1). Triangles, the best CTCF motif match (JASPAR MA0139.2): ▶, + strand; ◀, − strand. Positions in Mb (GRCh38). Orange arcs, pairs tethered in the model; grey dashed arcs with ×, pairs not tethered.

### Homologous copies are in closer contact in two duplicated loci where reads can be mapped

The PRAMEF locus cannot itself test whether homologous copies are closer in space, because its reads do not map (Fig 1B). We therefore turned to two duplicated loci whose paralogs have diverged enough for reads to map uniquely: the protocadherin cluster (chr5:140.70–141.60 Mb) and a zinc-finger cluster on chr19 (57.0–58.0 Mb). Fig 8A sets H1 Micro-C contact enrichment (above the diagonal) against repeat-masked sequence homology (below) for both clusters.

Contact enrichment compares how often two bins touch with how often any two bins the same distance apart touch. It is the observed/expected (O/E) value: the contacts of a bin pair divided by the average for all pairs at the same separation. A value of 1 means as often as expected, 2 means twice as often and 0.5 half as often. Distance alone raises contact, much as two houses on the same street meet more often than houses across town, so enrichment asks whether two pieces of DNA meet more often than their distance predicts. Repeat-masked sequence homology is the similarity between each pair of 2 kb bins after interspersed repeats were removed. These are short mobile elements, such as Alu and L1 sequences, present in thousands of copies across the whole genome. Left in, they would make almost any two bins look alike. Masking them is like ignoring common words such as “the” and “and” when comparing two texts: what remains are the copies made by local duplication (Methods).

We compared each homologous bin pair with a randomly chosen bin pair at the same genomic separation but without homology (Fig 8B). In both loci, homologous pairs had higher contact enrichment than their matched controls. The per-pair ratio divides the enrichment of each homologous pair by that of its control, so a ratio above 1 means the copies touch more often than unrelated DNA at the same distance. In the protocadherin cluster, the median observed/expected value was 1.14 for homologous pairs and 0.92 for controls (median per-pair ratio 1.21; 122 pairs; one-sided Mann–Whitney *P* = 1.4 × 10⁻⁴). In the zinc-finger cluster it was 1.66 and 0.96 (median ratio 1.86; 39 pairs; *P* = 1.0 × 10⁻⁴).

When the CSynth-like model folds a locus, each pair of homologous bins can be joined by a weak spring that pulls the two copies together. The strength of this spring is the homology weight, k. At k = 0 there is no pull, and the copies end up only as close as their distance along the chain allows. At larger k the copies are pulled closer, so the model shows more contact enrichment between them. We therefore asked how hard the spring must pull for the model to reproduce the enrichments measured by Micro-C (Fig 8C). We folded each homology map with a CSynth-like model at a range of homology weights *k* and measured the same enrichment in the folded paths. When only homologous pairs attract, enrichment rises with *k* and meets the Micro-C value at *k* = 0.0069 in the protocadherin cluster and 0.0071 in the zinc-finger cluster (Fig 8C, left). When the whole map attracts, the two loci give weights 3.5-fold apart (0.0039 and 0.0139). Matching the enrichment is a narrow test: a fold could bring the right copies together and still get the rest of the domain wrong. We therefore also compared the whole shape of each folded path with the path folded from the Micro-C map itself. Any two paths agree for bins that lie close along the chain, simply because neighbours stay near each other, so we removed this effect of genomic distance first and measured only the agreement beyond it. At no weight did the folded path agree with the path folded from Micro-C beyond what genomic distance alone predicts: the distance-corrected correlations ranged from −0.01 to 0.07, within one standard deviation of the values at *k* = 0 (Fig 8C, middle). Applied to the PRAMEF locus, the calibrated weight gives a 3.0-fold enrichment of homologous pairs, whereas the CSynth path, the refolded path and the final model are 7.0-, 9.7- and 8.3-fold enriched (Fig 8C, right). In short, one weak homology weight reproduces the Micro-C enrichment of both control loci, but it does not reproduce their overall fold, and the PRAMEF model holds its copies together about three times more tightly than this calibration supports.

**Fig 8.**
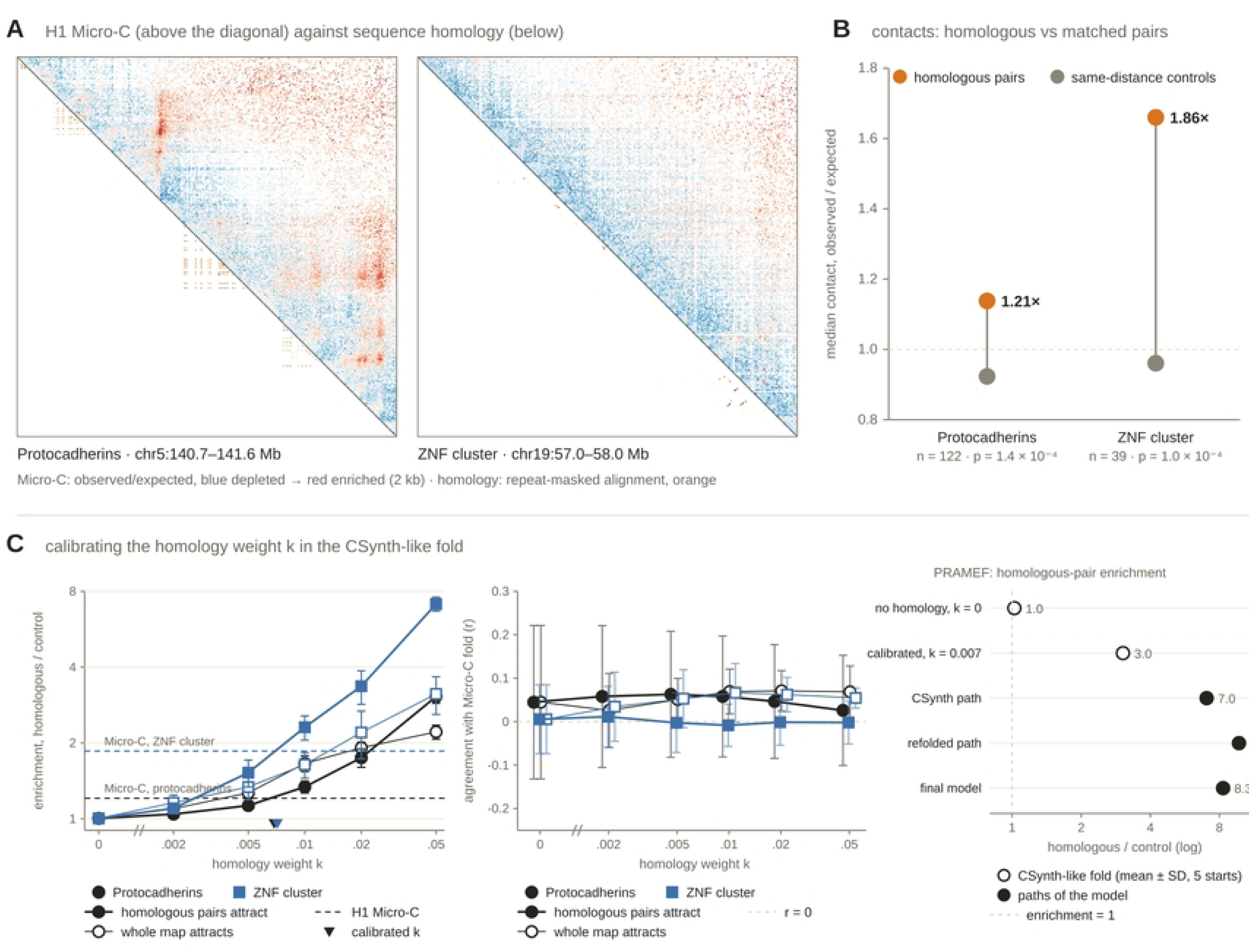
Micro-C contact enrichment and sequence homology in two duplicated loci with uniquely mappable reads. (A) The protocadherin cluster (chr5:140.7–141.6 Mb) and a zinc-finger (ZNF) cluster (chr19:57.0–58.0 Mb). Above the diagonal, H1 Micro-C contact enrichment (observed/expected, 2 kb bins; blue, depleted; red, enriched). Below the diagonal, repeat-masked sequence homology (orange). (B) Median contact enrichment (observed/expected) of homologous bin pairs (orange) and of same-distance controls (grey). For each homologous pair, one control was drawn at random from the bin pairs at the same genomic separation without homology (Methods). Vertical lines join the two medians for each locus; dashed line, observed/expected = 1. Labels give the median of the per-pair ratios (homologous/control), the number of homologous bin pairs (n) and the *P* value of a one-sided Mann–Whitney U test (homologous greater than control). (C) Calibration of the homology weight k in the CSynth-like fold (Methods). Left, contact enrichment of homologous over same-distance control bin pairs in the folded paths, with contact probability taken as proportional to distance⁻³, against k; k = 0 (no homology attraction) is plotted left of the axis break. Filled symbols and thick lines, attraction applied to homologous pairs only; open symbols and thin lines, attraction applied across the whole homology map; circles, protocadherin cluster; squares, ZNF cluster; error bars, SD over five random starts; dashed lines, the Micro-C enrichment of each locus from B; triangles, the weight at which the homologous-pairs-only curve meets the Micro-C value (linear interpolation). Middle, correlation between each folded path and the path folded from the H1 Micro-C map, after the log distances at each genomic separation were converted to z-scores; symbols as on the left, offset along k where they overlap. Right, homologous-pair enrichment in the PRAMEF locus for the CSynth-like fold at k = 0 and k = 0.007 (open circles; mean ± SD over five starts) and for the CSynth path, the refolded path and the path of the final model (filled circles); dashed line, enrichment = 1.

### Inside the PRAMEF locus, the two arms of a direct repeat are in contact in two cell types

At the scale of the whole locus, the few PRAMEF bins that survive balancing in Fig 1B carry no obvious structure. On closer examination, however, they hold one small pattern: a short line of contacts parallel to the diagonal, joining 13.225–13.245 Mb to the bins exactly 145 kb downstream (13.370–13.390 Mb; Fig 9A, arrowhead above the diagonal; enlarged in Fig 9B, arrow). The same pair of positions lies on the D3/D4 stripe of the SEQSIM map (Fig 9A, arrowhead below the diagonal). The line holds 291, 197, 129, 122 and 55 read pairs, against a mean of 14–32 for the bins beside it (Fig 9D). The 145 kb offset is the spacing between the two arms of the direct repeats D3 and D4. Rows where the two copies are 96–99% identical over 5 kb carry almost no uniquely mapped reads, so the line can be followed only over the 25 kb where the copies have diverged to 82–94% identity (Fig 9B, left).

A line at the copy offset is what two copies lying side by side would produce, but two other sources could give a similar signal. First, reads from one copy could be assigned to the other, moving contacts between neighbouring positions to the copy offset. This would also give a line, but one carrying only the small fraction of reads that are misassigned. Second, a structural variant in the H1 genome, such as a deletion or inversion with its junction between the two copies, would bring the sequences on either side of the junction together. A simple junction gives a single spot that fades away from one corner rather than a line, but a gene-conversion event that left a hybrid copy could give a line, so a variant specific to one genome had to be tested directly. We tested the first explanation by simulating reads from these bins and mapping them back to the locus (Methods). Reads of 50–150 bp were placed on the partner copy with mapping quality of at least 1 in at most 0.2% of cases, and never with mapping quality 30 or more. Applied to the contacts of each bin with its neighbours, this predicts at most 10 read pairs on the line in H1, against 55–291 observed (Fig 9D, crosses). To test the second, we turned to Micro-C from HFFc6 foreskin fibroblasts, a different individual and cell type [4]. The same line is present (Fig 9C): 456, 366, 115, 271 and 56 read pairs on the line, against 25–49 beside it and at most 19 expected from misassignment (Fig 9D). In both maps the line stands 2-to 10-fold above its neighbours, and the same bins at either end are discarded during balancing (Methods). The two arms of D3 and D4 are therefore in contact in two unrelated genomes, and this is the one measurement inside the PRAMEF locus that bears directly on the central assumption of the model, that homologous copies lie side by side. In the model, the two arms run side by side (Fig 9E). Nucleosomes 145.75 kb apart are 140–170 nm apart along the arms (13.185–13.220 Mb) and 95 nm apart at 13.225 Mb, where the Micro-C line begins, against a median of 471 nm for all positions 145.75 kb apart in the locus. Along the rest of the line (13.230–13.250 Mb) the two arms move apart again, to 140–280 nm (Fig 9E, orange).

**Fig 9.**
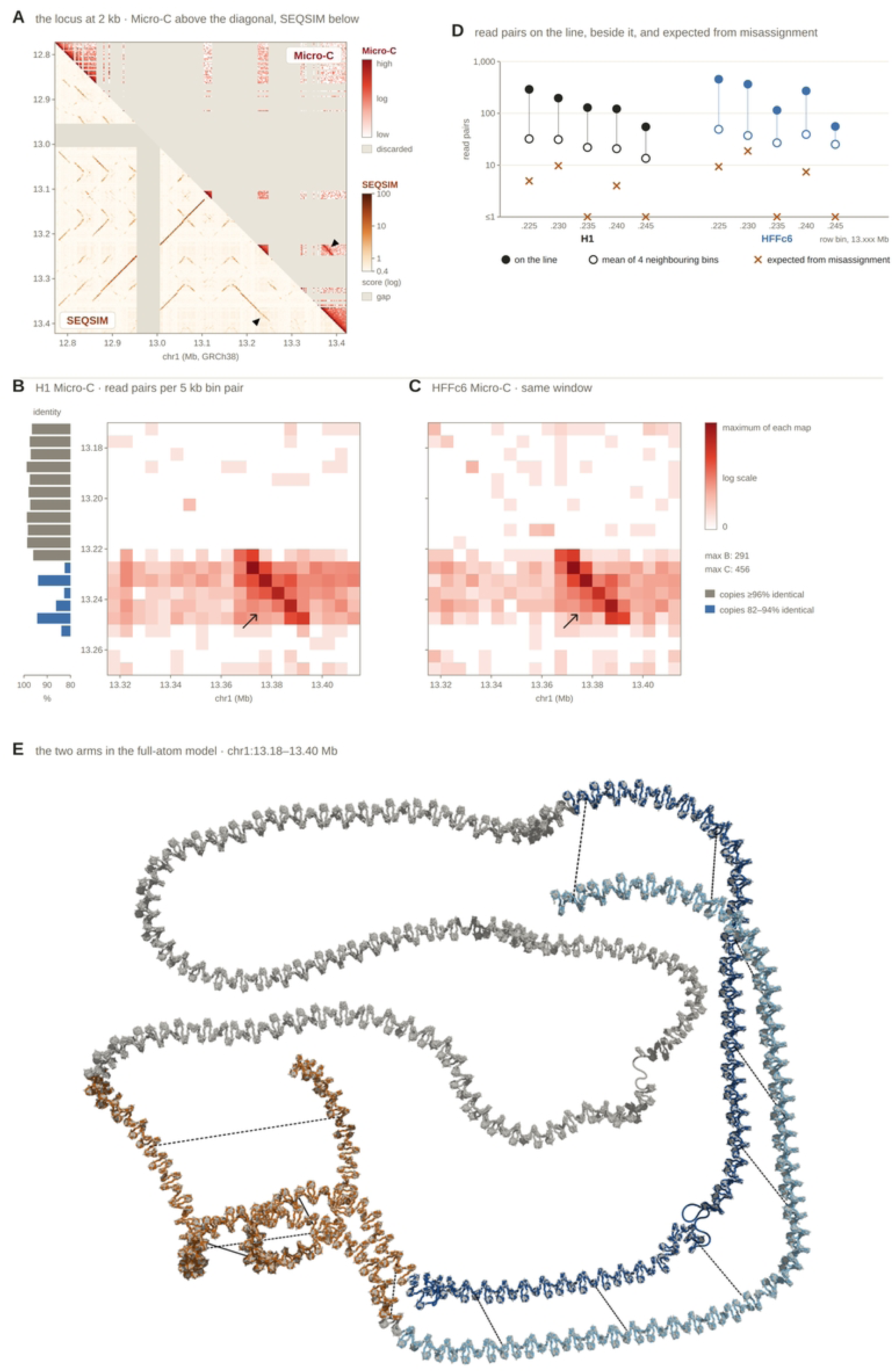
Micro-C contacts between the two arms of blocks D3 and D4 at a 145 kb offset in two cell types, with the counts expected from reads assigned to the wrong copy, and the two arms in the full-atom model. (A) The PRAMEF/HNRNPCL locus (chr1:12.77–13.42 Mb, GRCh38) in 2 kb bins. Above the diagonal, H1 Micro-C [4] (4DN 4DNFI9GMP2J8), balanced contacts on the logarithmic colour scale of Fig 1B; grey, bins discarded during balancing. Below the diagonal, the SEQSIM map of Fig 2A, logarithmic colour scale; grey, the assembly gap. Micro-C bins start 679 bp downstream of the SEQSIM bins. Arrowheads, contacts between 13.224–13.248 Mb and the bins 145 kb downstream (above the diagonal) and the same pair of positions in the SEQSIM map (below). (B) H1 Micro-C: read pairs per 5 kb bin pair before balancing. Rows, chr1:13.170–13.270 Mb; columns, 13.315–13.415 Mb, so that each column lies 145 kb downstream of the matching row and the diagonal from top left to bottom right joins positions 145 kb apart. Logarithmic colour scale from 0 to the maximum of the map. Arrow, contacts between 13.225–13.245 Mb and 13.370–13.390 Mb. Left, sequence identity between each 5 kb row bin and the bin 145 kb downstream (minimap2); grey, ≥96%; blue, 82–94%; blank, no alignment. (C) HFFc6 Micro-C (4DN 4DNFI643OYP9), same window and display as B. (D) Read pairs for each row bin from 13.225 to 13.245 Mb. Filled circles, the bin 145 kb downstream (on the line); open circles, mean of the four bins that share an edge with it; crosses, read pairs expected from reads assigned to the wrong copy, computed from simulated 50 bp reads with 0.5% sequencing error at mapping quality ≥1 and the contacts of each bin with its neighbours (Methods). Black, H1; blue, HFFc6. Logarithmic scale; values below 1 are plotted at 1. (E) chr1:13.18–13.40 Mb in the final full-atom model, DNA and histones drawn as lines. Light blue, 13.184–13.224 Mb (upstream arm of D3 and D4); dark blue, 13.330–13.370 Mb (downstream arm); orange, the 25 kb of each arm covered by the Micro-C line (13.225–13.250 and 13.370–13.395 Mb); grey, the rest of the window. Dashed lines join positions 145.75 kb apart, every 5 kb from 13.185 to 13.250 Mb. Thin loops, the long linkers at cohesin tether sites (Fig 7); rings not shown.

### MaaD-TAD builds all-atom models from measured Micro-C maps

To test the pipeline on an ordinary contact map, we built two loci that Micro-C covers almost completely, without homology restraints or tethers (Fig 10). IGHC (chr14:105,540,001– 105,750,000; 210 kb) holds the immunoglobulin heavy-chain constant genes, and PWRN (chr15:24,130,001–24,440,000; 310 kb) is a window on chromosome 15. Like PRAMEF, both contain duplicated segments, but H1 Micro-C discarded only 6 of 105 and 7 of 155 of their 2 kb bins, against 227 of 327 in PRAMEF. Each locus was folded from its H1 Micro-C map with CSynth and passed through Folder, Placer, Builder and Checker with the settings used for PRAMEF, giving models of 1,105 and 1,631 nucleosomes.

Fig 10A sets the H1 Micro-C map of each locus against its SEQSIM map. Both loci contain lines of sequence similarity, most of them parallel to the diagonal, which mark duplicated segments in the same orientation. In IGHC the longest line joins the region around IGHA2 and IGHE to the region around IGHA1 and IGHG1, the same two regions joined by the strongest line of the Micro-C map. In PWRN the longest line is L4, whose two arms are marked above the map. With the test of Fig 8B, homologous bin pairs had 8.8 times (IGHC; 123 pairs) and 4.6 times (PWRN; 139 pairs) the H1 Micro-C contacts of control pairs at the same separation (one-sided Mann– Whitney p ≤ 3 × 10⁻²⁴; HFFc6, 11.9 and 3.3 times). These values are not an independent estimate of the enrichment in Fig 8: both windows were chosen from a scan of the H1 map for lines of contacts parallel to the diagonal, the kind of line this test rewards, so they overstate it. A genome-wide measure is in preparation. These lines were not used to build the models in Fig 10B–D, which were folded from Micro-C alone.

Fig 10B sets the contacts of each model against the Micro-C map it was built from. The strongest line of the IGHC map, which joins the region around IGHA2 and IGHE to the region around IGHA1 and IGHG1, appears in the model close to its position in the map (Fig 10B, left). The models, built without homology terms, kept part of the enrichment of Fig 10A: homologous pairs had 3.0 (IGHC) and 2.4 times (PWRN) the model contacts of control pairs at the same separation (p ≤ 4 × 10⁻¹⁵). To compare the two triangles over the whole map, we removed the dependence of contacts on genomic distance and correlated model and map within each separation. On 10 kb blocks, the models agreed with H1 Micro-C with r = 0.61 (IGHC) and 0.39 (PWRN). This matches the CSynth folds they were built from (0.63 and 0.38) and approaches the agreement between the H1 and HFFc6 maps of the same loci (0.74 and 0.33), which sets the ceiling for any single structure. At 2 kb all of these values were low (models 0.13 and 0.09; H1 against HFFc6, 0.16 and 0.05). Through the pipeline, the full-atom models kept the fold of their CSynth paths (r = 0.81 and 0.85 between the distance patterns).

Fig 10C shows the two models whole, at the same scale. IGHC is 1.19 µm across at its largest extent (radius of gyration 414 nm) and PWRN 1.62 µm (561 nm). Fig 10D shows eight consecutive nucleosomes of each model at atomic detail, within IGHG1 and within arm 1 of L4, the longest line of the PWRN SEQSIM map.

Both models passed every Checker test. All 210,000 and 310,000 base pairs are present on both strands, and none of the 209,999 and 309,999 O3′–P linkages per strand is broken, including those across 22 and 32 block seams. At most 28 phosphodiester linkages per model are strained (0.01%), and there are no long-range clashes. On the workstation used for PRAMEF, building took 21 and 34 min, and tail repair 10 and 27 min.

**Fig 10.**
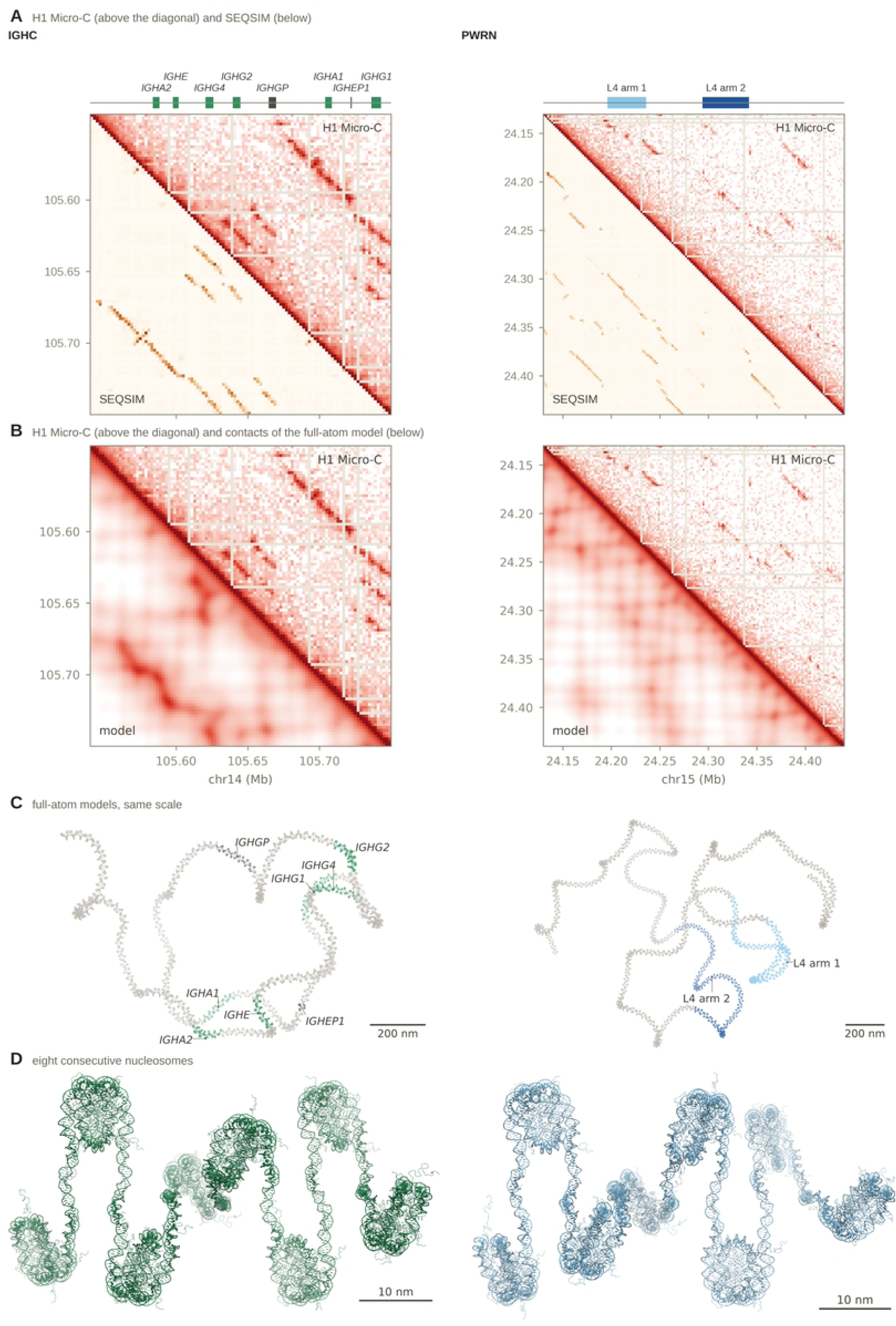
All-atom models of two loci built from Micro-C alone. (A) Contact maps of IGHC (chr14:105,540,001–105,750,000; left) and PWRN (chr15:24,130,001–24,440,000; right) in 2 kb bins. Above the diagonal, H1 Micro-C [4] (4DN 4DNFI9GMP2J8; balanced counts) on a log scale, clipped at the 5th and 99.3rd percentiles, with the colour ramp of Fig 1B; light grey rows and columns, bins discarded by the 4DN pipeline. Below the diagonal, the SEQSIM map of the same window (the larger of the direct and inverted scores of each pair of bins) on a log scale from 0.4 to 100, with the SEQSIM colour ramp of Fig 9A. Tracks above the maps: IGHC, immunoglobulin heavy-chain constant genes (green) and pseudogenes (dark grey) (GRCh38, Ensembl); PWRN, the two arms of L4, the longest line of the PWRN SEQSIM map (26 bin pairs with SEQSIM > 1.1; arm 1 light blue, arm 2 dark blue). (B) The same Micro-C maps (above the diagonal) against contacts of the full-atom model (below), computed as distance⁻³ between the mean nucleosome centres of each pair of bins, on a log scale clipped at the 5th and 99.3rd percentiles of their own values, with the colour ramp of Fig 1B. Genomic coordinates as in A. (C) The full-atom models, DNA and histones drawn as lines (grey), with the genes and arms of A coloured as in A. Each model is viewed along the two longest principal axes of its nucleosome centres; depth cueing on. Both models are drawn at the same scale. Scale bar, 200 nm. (D) Eight consecutive nucleosomes of each model, cartoon, coloured as in C: IGHC, nucleosomes 1,046– 1,053 in IGHG1 (dyads chr14:105,738,902–105,740,232); PWRN, nucleosomes 448–455 in L4 arm 1 (dyads chr15:24,215,236–24,216,566). Scale bar, 10 nm.

## Discussion

We have built an all-atom model of a 652 kb chromatin domain in a region where both contact maps and nucleosome maps are largely missing (Fig 1). The model places 652,000 bp of DNA on 3,359 nucleosomes, with every linker closed at atomic detail and no DNA or histone passing through another part of the model (Fig 5, Table 3). Four steps made this possible: sequence homology in place of measured contacts (Fig 2), refolding of the homology-derived path so that it can carry a chromatin fibre (Fig 3), restraints that hold homologous copies in register (Fig 6), and cohesin tethers between CTCF sites and their inverted copies (Fig 7). The model is therefore both a test of the pipeline under difficult conditions and an explicit, testable hypothesis about how a duplicated domain could be folded.

### From a contact map to a fibre

A path folded from a contact map indicates which loci are near each other, but not the fibre that joins them. Scaled to a domain of realistic size, the CSynth path was 8.5 times too short to hold the nucleosomes of the locus laid end to end, and 2.6 times too short to hold them as a compact two-start fibre. A third of it came closer to another part than the fibre is wide, and 18 bends were tighter than the fibre can turn (Fig 3). These problems are not specific to CSynth or to this locus. Any model in which one bead stands for kilobases of DNA must be lengthened, and given thickness and stiffness, before nucleosomes can be placed on it. The refolding kept the overall fold (r = 0.91 between pairwise distances) while the domain grew from 0.5 µm, the scale imposed on the CSynth path, to 0.96 µm. The fold encoded in the SEQSIM map survived conversion into a buildable fibre, and we expect the same of measured contact maps, which share its kilobase-scale beads.

The fibre we built is a regular two-start zigzag, the arrangement seen in the crystal structure of a tetranucleosome [36]. In nuclei, chromatin appears instead as a flexible, disordered chain 5–24 nm in diameter, packed at varying density [37]. The 52 nm spacing and 123 nm minimum bend radius imposed here are conservative choices that make a clash-free build possible, not measurements. A less regular fibre would pack the same DNA into a smaller volume, and the pipeline can accept such geometries once they are defined for a region.

### Homology as a proxy for proximity

The central assumption of the model is that homologous copies lie closer together than random pairs at the same genomic distance. Over most of the PRAMEF locus this cannot be tested, because its reads do not map (Fig 1B). In two duplicated loci whose paralogs have diverged enough for reads to map, homologous bin pairs had 1.21- and 1.86-fold higher Micro-C contact enrichment than distance-matched pairs without homology (Fig 8B). This agrees with genome-wide observations that paralogous genes contact each other more often than distance-matched gene pairs and share enhancers more often [12], and that repeats of the same family cluster in the nucleus [15].

The enrichment is consistent but modest, and the control loci differ from PRAMEF in one important respect: their copies are fewer and less often near-identical. Aligned the same way, the protocadherin and zinc-finger windows hold 133 and 41 homologous 2 kb bin pairs, against 829 in PRAMEF, and no protocadherin pair and 10% of zinc-finger pairs exceed 97.9% identity, against 22% in PRAMEF (S3 Table). Both are also older gene families: the protocadherin clusters date to early vertebrates [38], and the 19q13.43 zinc-finger clusters have orthologs in dog and mouse [39], whereas the near-identical PRAMEF copies arose by a more recent expansion [16]. Whether near-identical copies such as those of PRAMEF are held closer than diverged ones cannot be decided from these data. The calibration in Fig 8C puts a number on this. One homology weight reproduces the Micro-C enrichment in both control loci, and when applied to PRAMEF, it predicts a 3.0-fold enrichment of homologous pairs, whereas the final model is 8.3-fold enriched. The model therefore sits closer to the strong-homology limit than the control loci support and should be read as the fold expected if homology is the dominant signal. The calibrated term also did not reproduce the rest of the Micro-C fold in either control locus: homology predicts which pairs come together, not the overall shape of the domain. Because the PRAMEF copies are younger and more alike than those of the control loci, they may pair more strongly than this calibration implies; the 145 kb line between D3 and D4 (Fig 9) is one place where that strength could be measured directly. A genome-wide survey of SEQSIM patterns on every chromosome (manuscript in preparation) extends this comparison beyond two control loci and asks which kinds of duplication SEQSIM predicts best.

Inside the PRAMEF locus itself, one measurement bears on this directly. Where the arms of D3 and D4 have diverged enough for reads to map, H1 Micro-C shows a short line of contacts at exactly the 145 kb offset between them, parallel to the diagonal (Fig 9A and 9B). This is the pattern expected if the two direct copies lie side by side in register, as they do in the model (Fig 6). Reads assigned to the wrong copy explain at most a few percent of these contacts (Fig 9D). Two other explanations remain. A deletion or duplication between the repeats in H1 cells would concentrate contacts near a single junction rather than spread them along a line, but a hybrid copy formed by gene conversion, carrying stretches of both arms, could produce a line. Such variants differ between individuals, yet the line is equally clear in Micro-C from HFFc6 fibroblasts, a different person and cell type (Fig 9C). This argues against a variant unique to H1 cells. A variant common in the population, or an error in the reference assembly shared by both genomes, would still produce the line in both maps. Remapping to the complete telomere-to-telomere assembly of the human genome (T2T-CHM13) [40], or long-read contact maps that resolve the copies directly, would close this gap (see Predictions). If the line reflects pairing, it is the one direct measurement inside the locus that supports the central assumption of the model.

The model itself matches it only in part: it brings the arms closest where the line begins (95 nm at 13.225 Mb) and lets them drift apart along the rest of the line (140–280 nm; Fig 9E). It therefore reproduces the side-by-side arrangement of the arms but not the contacts along the diverged 25 kb. Homology attraction is weakest there, because the copies are only 82–94% identical, which may be why the model loses the contact where Micro-C detects it.

### Register and mirror-image genes

Contact maps tell which copies meet but not how they align. The CSynth path brought homologous arms together but left them out of register by about ten nucleosomes (median 2,018 bp), and refolding alone did not improve this (Fig 6B). Pairing restraints reduced the error fifteen-fold, to less than one nucleosome repeat, at a moderate cost to the original fold (r from 0.91 to 0.80). In register, mirror-image genes on opposite arms of the I11/I12 hairpin lie opposite one another, with HNRNPCL2 at the tip (Fig 6C). If copies are aligned this way in cells, the promoters of each mirror pair would share a local environment, the arrangement proposed in the Introduction (premise 3). Aligned near-identical copies are also the substrates of gene conversion and non-allelic recombination, so the same geometry could contribute to the rearrangements that recur in duplicated regions.

Block D1 shows a limit of pairwise restraints. Its first arm is also an arm of I1 and I2, and one stretch of fibre cannot be in register with three partners at once. In a population of cells such a stretch may alternate between partners, which a single static model cannot represent.

### Copied CTCF sites as loop anchors

Inverting a duplicated segment necessarily turns a copied CTCF site into either a convergent or a divergent pair with its source. Convergent CTCF sites are the preferred anchors of cohesin-mediated loops, and inverting a single site can remove or reroute a loop [1–3]. Loop extrusion explains this preference: cohesin extrudes DNA until it meets CTCF bound in the orientation that faces it [41–43]. Of the five inverted pairs with ChIP-seq peaks at both copies, the three we tethered are convergent and the two left untethered are divergent (Fig 7D). Because the tethers were placed by us, the model does not show that these sites anchor loops in cells; it shows that the orientation of the copied sites is compatible with that role.

Fig 11 summarizes the model that links these observations. A duplication copies a segment, including its CTCF site, to a part of the fibre that is already nearby (Fig 11A). An inverted copy faces its source (Fig 11B). Cohesin extruding the loop between them stops at the convergent pair (Fig 11C), and the aligned copies, with their promoters, share a microenvironment (Fig 11D). In this view, duplication alone can create a loop anchor without any new binding site. The atomic model shows that the geometry is at least sterically possible: both naked duplexes pass together through a ring with an opening only 7 nm across, far smaller than the roughly 40 nm ring of cohesin [44], with no other DNA or histone inside it and no atom closer than 1 nm to the ring (Fig 7C).

**Fig 11.**
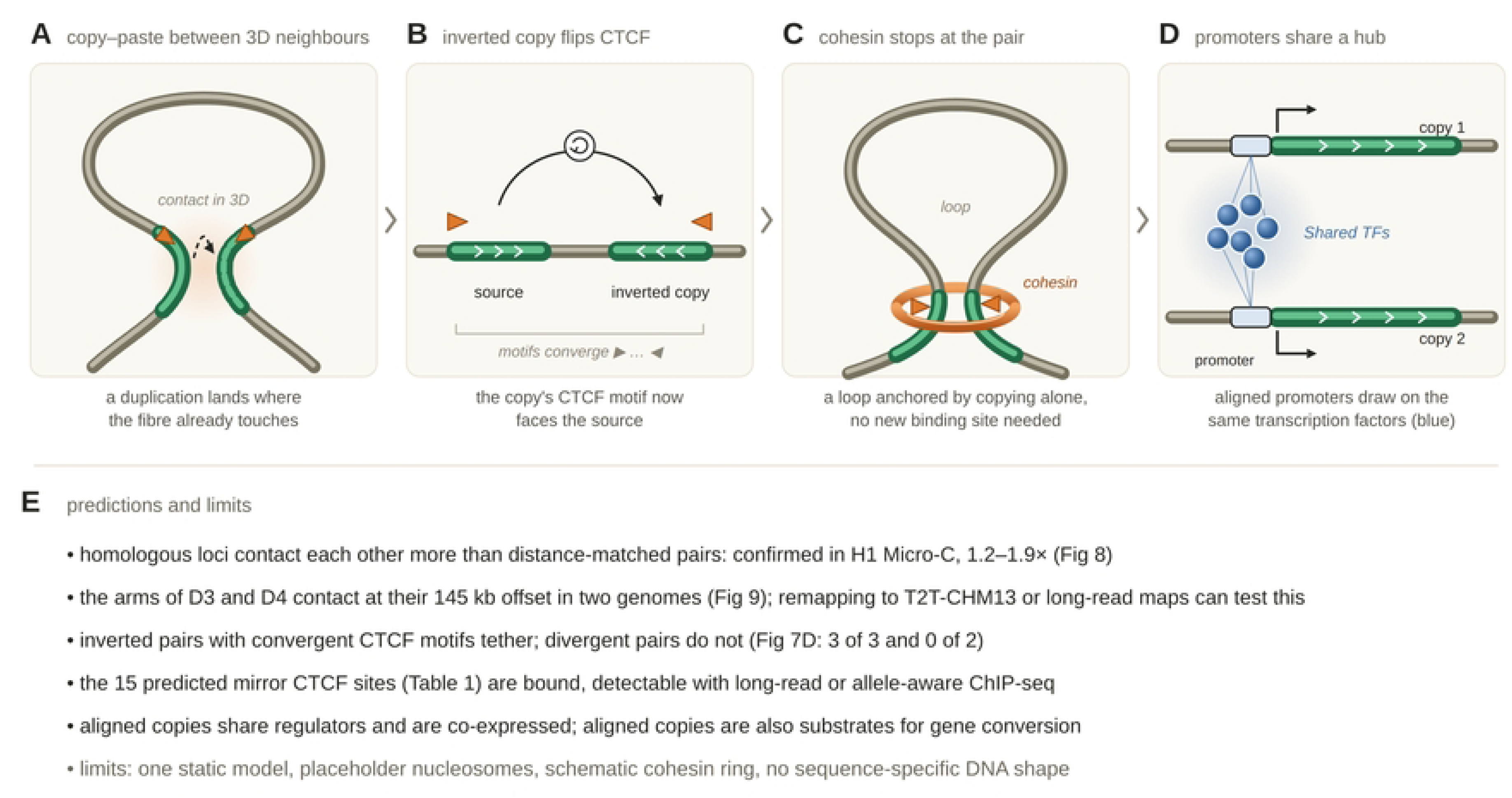
Schematic model linking segmental duplication, CTCF motif orientation and spatial proximity. (A) A duplication (dashed arrow) copies a segment (green) carrying a CTCF site (orange triangle) to a part of the fibre (grey) that is close in three dimensions (shaded); the new copy has a dashed outline. (B) The source and its inverted copy along the chromosome; white chevrons, orientation of each copy; orange triangles, CTCF motifs (▶, + strand; ◀, − strand). (C) A cohesin ring (orange) holding the loop at the two convergent sites. (D) Two aligned copies (green bars), each with its promoter (outlined box, upstream of the copy) and transcription start (bent arrow); blue circles, transcription factors (TFs) shared by the two promoters, with lines joining each TF to both promoters. (E) Predictions and limitations of the model, with the figures that address them. All panels are schematic and not to scale.

### What the model suggests for the PRAMEF/HNRNPCL locus

DUX4 switches on PRAMEF genes at the cleavage stage [19] and activates HNRNPCL1 [20]. The model places mirror-image genes side by side and in register, with their promoters close together (Fig 6C). If the copies are paired in the nucleus, a regulatory region bound by DUX4 could serve both copies of a pair, helping a short burst of transcription reach every copy at once. This can be tested directly, by measuring the distance between the promoters of mirror-image genes in cells that model the cleavage-stage embryo, and by deleting or inverting a tethered CTCF site (Predictions). The same DUX4 targets are switched on in the wrong tissue in FSHD [20], so the fold of the locus may also bear on disease. Finally, because the locus changed length between reference versions and still contains a 50 kb gap, the model gives a frame in which missing sequence and differences between people can be placed, and their effect on contacts between genes predicted.

### Predictions

The model makes predictions that do not depend on mapping short reads. First, the fifteen predicted CTCF sites (Table 1), including six copies of cohesin sites, should be bound wherever their ChIP-supported copies are bound. Long-read methods that record protein–DNA contacts on single molecules, such as DiMeLo-seq [45], can place reads within near-identical copies and test this directly. Second, the complete T2T-CHM13 assembly [40] has no gaps, including the 50 kb gap that remains in this locus in GRCh38; long-read or multi-contact maps anchored in that assembly could test the predicted pairing of mirror-image genes. Third, deleting or inverting one copy of a tethered pair should loosen the corresponding hairpin, which can be tested by imaging the distance between unique sequences flanking the two copies.

### Limitations

The model has several limitations. Eighty-five percent of nucleosome positions are placeholders at a consensus spacing, and the measured 15% come from one part of the locus. The fibre is a regular zigzag without linker histone, histone variants or modifications. The model is a single static structure, whereas chromatin in cells is an ensemble of conformations that differs between cells and over time. Cohesin is represented by a schematic ring, and the tethers were placed rather than predicted. The structure has not been relaxed with a molecular force field, so its local geometry is that of the building blocks and the closure solver rather than of an equilibrated structure. About 68,000 atom pairs between neighbouring nucleotides lie closer than 0.22 nm, local strain that a brief energy minimization would be expected to remove, and the histone tails were moved only far enough to clear the DNA, not sampled. The sequence threaded onto the model is that of the GRCh38 reference, which is assembled from several donors rather than from one person’s genome. Individuals may carry more or fewer copies of a duplicated locus, so the reference sequence may not represent the locus in any given cell; indeed, the copy number of PRAMEF1 differs between human populations [7]. Finally, the 50 kb assembly gap was filled without sequence information, so the path through it is a smooth placeholder arc and the contacts it makes carry no predictive weight.

### Conclusions

We created MaaD-TAD, a model builder that generates all-atom models of complete TADs from publicly available data, and that can build models where those data have gaps by stating explicitly each assumption it makes. Applied to the PRAMEF/HNRNPCL locus, it produced 652 kb of chromatin in which no DNA or histone overlaps another part of the structure, together with predictions that Hi-C and nucleosome mapping could not provide, such as CTCF binding at unmappable sites and the side-by-side positioning of near-identical genes, which can be tested with long-read methods.

Because each stage of MaaD-TAD uses standard inputs (a contact map, nucleosome positions and a list of tethers), it can also be applied to domains where measured data are available. There it can provide atomic starting structures for simulation, and for multidimensional studies that overlay genomic data on the structure.

If homology does mark proximity, as the Micro-C tests here suggest for the loci examined, MaaD-TAD offers a way to model chromatin structure even in regions that mapping data cannot reach.

## Supporting information

**S1 File. MaaD-TAD code and figure scripts.** The four MaaD-TAD (Molecular all-atom Display of TADs) modules, Folder (refolding), Placer (nucleosome placement), Builder (all-atom construction) and Checker (quality control and repair), and the scripts and input files that regenerate the figures (folder figures). The builds from Micro-C alone (Fig 10) are in folder hic_test (extract_hic_test.py, to_contacts.py, build_locus.py, thread_locus.py, score_models.py, homology_test_loci.py, and the figure scripts fig_seqsim_maps.py, fig_models_maps.py and make_figS_models_pml.py); S1 Video is rendered by s1_video.pml (folder pymol); the identity comparison of homologous copies is control_identity.py (folder control_loci). The same files are at https://github.com/mmarcet-palacios/MaaD-TAD, archived at Zenodo (https://doi.org/10.5281/zenodo.23718187).

**S1 Table. Key terms used in this paper.** Each term with a short plain-language explanation.

**S2 Table. Parameters of the refolding, pairing and tether restraints.** Values used to build the final model, grouped by MaaD-TAD module, with the script that applies each one (S1 File). Distances are in nm; weights are relative weights of the energy terms and have no units.

**S3 Table. Identity of homologous 2 kb bin pairs in PRAMEF and the two control loci.** Windows: PRAMEF/HNRNPCL, chr1:12,771,321–13,421,320 (325 bins); protocadherin cluster, chr5:140,700,001–141,600,000 (450 bins); zinc-finger cluster, chr19:57,000,001–58,000,000 (500 bins); GRCh38. Repeats and N were removed, bins with fewer than 300 unmasked bases were excluded, and every pair of bins was aligned on both strands (Smith–Waterman, parasail; match +1, mismatch −3, gap opening 5, gap extension 2). Homologous pairs are those scoring ≥ 30 (alignment score divided by the length of the shorter unmasked bin, × 100), as in Materials and methods; the protocadherin and zinc-finger counts reproduce the 133 and 41 pairs reported there. Identity, matching bases over the aligned columns of the traceback, gaps included, with unaligned flanks excluded. IQR, interquartile range. Script: control_identity.py (S1 File).

**S1 Fig. Nucleosome placement: steering and genomic anchoring.** (A) The 3,359 nucleosome centres of the model, one point each, in the orientation of Fig 4A; blue, nucleosomes with a measured dyad; grey, placeholders. Scale bar, 100 nm. (B) Distribution of the steering rotation applied to each nucleosome to follow the refolded path (0.5° bins); dashed lines, median and 95th percentile; nucleosomes above 20° are counted but not plotted. (C) Offset along the path between each nucleosome and its genomic position, against position in the locus, for placement anchored to the genome (green, solid) and for the same placement without anchoring (rust, dashed); positive, ahead of its genomic position; dashed grey line, zero offset. The labels give the median and 95th percentile of the absolute offset. The offsets of the last few nucleosomes (−140 nm in both runs) lie off scale (arrowhead).

**S2 Fig. Domain size of the CSynth path: reachability and sterics.** The CSynth path (326 beads of 2 kb) was scaled to domain diameters of 400–1,600 nm (largest distance between beads outside the assembly gap), and each 2 kb section was given the fibre axis provided by its nucleosomes (3,376 nucleosomes on a two-start zigzag; 23.5 µm of axis in total) (scale_feasibility.py, S1 File). (A) Number of sections, of 325, that are longer on the scaled path than the fibre axis their nucleosomes provide, so that the fibre cannot span them. Blue band, domains at which every section is reachable. (B) Radius of the helical coil needed to fold the extra fibre into each section (orange circles) and the largest coil radius that keeps the fibre clear of other parts of the path, given the path’s closest self-approach and a fibre radius of 11 nm (blue squares). Orange band, domains at which the coil fits. Vertical dashed line, 500 nm, the scale used for the CSynth path (Methods). Boundaries of the bands are interpolated between the scanned diameters.

**S3 Fig. Backfolding as an alternative to a helical coil.** Instead of coiling, the extra fibre of each 2 kb section of the CSynth path was folded back on itself in a flat serpentine of arcs at the fibre’s minimum bend radius (49.9 nm, set by a steering budget of 8° per nucleosome) and straight legs, with the flat side of each fold facing the nearest neighbouring strands (backfold_axis.py, S1 File). The path was scaled to domain diameters of 500–1,100 nm. (A) Smallest distance between fibre axes of non-adjacent parts of the path; dashed line, 22 nm, two fibre radii. (B) Number of 2 kb sections, of 325, with a fibre clash. (C) Median relative difference between the fibre length delivered to each section and the length its nucleosomes require. (D) 95th percentile of the bending required per nucleosome step; dashed line, the 8° per nucleosome that linkers can absorb. Points are the scanned diameters joined by lines.

**S1 Video. The full-atom model of the PRAMEF/HNRNPCL locus, turned through 360°.** The model of Fig 5, in the orientation of Fig 5A, rotated once about the vertical axis of the image (0.4° per frame; 900 frames at 30 frames per second, 30 s). All atoms drawn as lines; DNA blue (RGB 0.247, 0.435, 0.659), histones orange (RGB 0.851, 0.451, 0.122); white background; orthographic projection with depth cueing (distant parts fade toward white). Rendered in PyMOL draw mode at 1,920 × 1,080 pixels and encoded as H.264 (s1_video.pml, S1 File).

**S1 Text. Figure preparation.** How each figure panel was drawn, so that it can be regenerated from the scripts in S1 File.

